# A systematic review and meta-analysis of genetic parameters for complex quantitative traits in aquatic animal species

**DOI:** 10.1101/2021.05.20.445048

**Authors:** Nguyen Hong Nguyen

**Author notes:** Correspondence to: Dr Nguyen Hong Nguyen.

## Abstract

A systematic review and meta-analysis of genetic parameters underlying inheritance and complex biological relationships for quantitative traits are not available for aquatic animal species. I synthesised and conducted a comprehensive meta-analysis of the published information from 1985 to 2017 on heritability, common full-sib effects and genetic correlations for quantitative characters of biological importance (growth, carcass and flesh quality, disease resistance, deformity and reproduction) for aquaculture species. A majority of the studies (73.5%) focussed on growth related traits (body weight), followed by those on disease resistance (15.9%), whereas only a limited number of studies (10.6%) reported heritability estimates for carcass and flesh quality, deformity or reproduction characteristics. The weighted means of heritability for growth (weight, food utilisation efficiency, maturity) and carcass (fillet weight and yield) traits were moderate. Resistance against various bacteria, virus and parasites were moderately to highly heritable. Across aquatic animal species, the weighted heritability for a range of deformity measures and reproductive traits (fecundity, early survival) was low and not significantly different from zero. The common full-sibs (c^2^) accounted for a large proportion of total variance for body traits but it was of smaller magnitude in later phase of the growth development. The c^2^ effects however were not significant or in many cases they were not reported for carcass and flesh quality attributes as well as survival and deformity. The maternal genetic effects were not available for all traits studied especially for reproductive and early growth characters. Genetic correlations between body and carcass traits were high and positive, suggesting that selection for rapid growth can improve fillet weight, a carcass trait of paramount importance. Body weight, the most commonly used selection criterion in aquatic animals, showed non-significant genetic correlation with disease resistance, likely because both positive and negative genetic associations between the two types of traits. Interestingly the genetic associations between growth and reproductive performance (fecundity) and fry traits (fry weight, fry survival) were favourable. To date, there are still no published data on genetic relationships of carcass and flesh quality with disease resistance or reproductive performance in any aquaculture species. Additionally, the present study discussed new traits, including functional, immunological, behavioural and social interaction as well as uniformity that are emerging as potential selection criteria and which can be exploited in future genetic improvement programs for aquatic animals.

## I. INTRODUCTION

Quantitative genetic parameters (heritability, maternal and common full-sib effects, phenotypic and genetic correlations) provide information needed for development of breeding objectives, design of selection strategies as well as assessment of selective breeding genetic improvement programs. Across aquatic animal species, heritability and trait correlations are often estimated in experimental populations prior to selection in order to examine genetic characteristics of new traits before closed nucleus breeding populations are initiated (Nguyen 2016; Olesen et al. 2003). As part of the process of monitoring selective breeding programs, the genetic parameters are re-estimated to ensure the continued response of traits under selection (i.e., selection criteria) and to predict possible changes in correlated characters to refine the breeding strategy. A large number of studies have reported heritabilities, maternal and common environmental effects and genetic correlations for traits of economic importance (e.g., body weight and survival) in important aquaculture species such as salmonids (Atlantic salmon, rainbow trout), carps, tilapias, shrimps, molluscs and newly cultured species (section 2.1 below). By contrast, genetic inheritance of flesh quality, behaviour and fitness-related traits is not well documented, although a few recent studies shown heritable additive genetic variations in sexual maturity, deformity and survival (Thoa et al. 2016), social interactions, and behavioural response characteristics (Drangsholt et al. 2014). Genetic correlations between growth performance and traits of commercial importance, such as flesh and eating quality, reproduction, fitness and functional adaptation traits, are not well understood. It is thus not possible to make any prediction of correlated changes in these traits to selection for high growth, a trait of paramount importance in almost all genetic enhancement programs for aquaculture species. On the other hand, measuring realised correlated response in these traits to selection is costly and time-consuming and an issue that remains unresolved because it requires multigenerational in-depth pedigree data to assess the changes with confidence (Hamzah et al. 2014b; Nguyen et al. 2010a). The possible changes on one trait to selection for another can be predicted from the knowledge of genetic and phenotypic correlation estimates.

While there is a paucity of published information in the literature to aid our understanding of genetic inheritance for commercial traits, especially those that are difficult to measure, a growing number of attempts has been made to develop systematic breeding programs for new species, including marine finfishes (Knibb et al. 2016) or spiny rock lobsters of high economic value (Nguyen et al. 2018a). As a first step to start genetic breeding programs for a new species, simple questions that breeders often ask is which traits they should select for, whether these traits are heritable and can respond to selection, and whether there are any consequences of selection on other characters of commercial interest. When the breeding program proceeds, expansion of the breeding goals also requires accurate estimates of genetic parameters for candidate traits. To obtain a reliable set of heritability and genetic correlations for new traits, rigorous experimental designs with reasonable family structure and sample size (Lynch and Walsh 1998) should be used. Taking these issues collectively, there is a need for a systematic review and meta-analysis of the published literature to guide decisions on future design and development of genetic improvement programs for aquatic animal species. Such a study has not been conducted for aquatic species, and weighted values of heritability and estimated correlations for commercial characters are currently not available.

The aim of this study, thus, was to synthesise information about inheritance and relationships among quantitative traits of economic importance in aquaculture species. Further, I also discussed the experimental design (i.e., sample size) required to obtain reliable genetic parameter estimates as well as proposing future directions to dissect genetic architecture of complex traits, especially novel characters that are emerging as potential selection criteria in selective breeding programs for aquaculture species.

## II. MATERIALS AND METHODS

### 1. Search strategy, selection criteria and traits

A systematic search of public electronic databases (PubMed, ISI Web of Knowledge, CABI and Google Scholar) using five keywords: “heritability, genetic parameters, correlations, variance and covariance components and aquaculture species” was undertaken to identify studies published in international, peer-reviewed journals from 1985 to 1^st^ March 2017. Published work prior to 1985 on genetic variation in quantitative traits in fish and shellfish was reviewed by Gjedrem (1983) and Gjerde (1986). In this study, a total of 1,120 records was identified from the databases. They were selected for inclusion in this review following the PRISMA (the Preferred Reporting Items for Systematic reviews and Meta-Analyses) protocol (Moher et al. 2009) (Supplementary file 1). Initially, their abstracts were screened, 260 duplicate or irrelevant records (i.e., papers on human, plant or terrestrial farmed animals or those did not report genetic parameter estimates) were excluded. The remaining 860 articles were examined for their full text, and 304 records were further excluded on the basis of the following criteria: 1) studies with less than 15 families due to their low statistical power to obtain reliable genetic parameter estimates (see Section 2.4), 2) reports with negative heritability or genetic and phenotypic correlations out of parameter space (−1 to 1), 3) studies published in languages other than English, 4) obviously irrelevant literature (i.e., papers on aquatic animals other than aquaculture species or those that reported traits that were not examined in this review), and 5) non-peer reviewed materials or project reports. After the assessment, 556 articles were assessed by studying their full texts and only 300 articles were eligible for inclusion in the review (256 references were discarded, mainly due to 1-5 as above). Our final screening also removed eight articles. A final list of 292 articles was used in our meta-analysis (full list of the references is given in Supplementary file No.2).

The dataset was then extracted from these (292) papers that included name of authors (references), Latin name of species, common species name, data records (observations), numbers of sires and dams (family), mating design, analytic methods, age of the animal, basic statistics for traits (mean, standard deviation and coefficient of variation), genetic parameters (heritability, common full-sib effects, phenotypic and genetic correlations) and their associated standard errors reported for traits of commercial importance.

Traits were grouped in six broad categories: 1) growth-related traits (body weight and feed utilisation efficiency), 2) carcass (fillet weight and fillet yield) and flesh quality (pH, colour, fat content) traits, 3) reproductive characteristics (sexual maturity, female weight before or after spawning, fecundity, fry weight, fry survival), 4) diseases resistance (virus, bacteria, parasite or others), and 5) fitness traits, mainly morphological deformity and survival during grow-out.

### 2. Meta-analysis

Firstly, I derived standard errors (*s*) for heritability and common full-sib effects for studies in which they were not reported. The theoretical *s* can be calculated based on the heritability estimate, number of offspring per sire and number of sires (Falconer and Mackay 1996). It can be also calculated approximately as 32h^2^/*n* where *n* is the number of data records (Koots et al. 1994). Alternatively, the SE of the heritability and common full-sib estimates for these studies were calculated by dividing weighted mean standard deviation (SDw) by the square root of the number of records in the estimate, using Equation 1 (Borenstein et al. 2011).

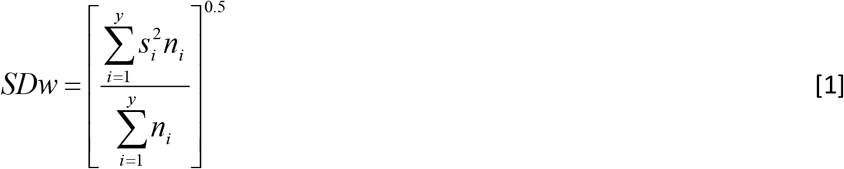

where *s* is the standard error and *n* is the number of records for the *i*^th^ estimate (*i*= 1,2,3…*y*).

Correlations among the three methods were greater than 0.9; hence only results based on the number of data records (Equation 1) are used in this study. I then weighted each estimate of a parameter from studies by their sample size or the inverse of variance. Both of the weighting methods gave similar weighted mean heritability for growth or reproductive traits, but the standard errors from the latter method (the inverse of the variance) were smaller than those from the former. Therefore, weighted means for basic statistics (trait mean, SD, CV) for all traits were derived using sample size as weight. On the other hand, weighted mean heritability and common full-sib effects were obtained using the inverse of the variance as weight (Akanno et al. 2013; Safari et al. 2005).

To calculate the weighted mean estimate of phenotypic and genetic correlations between traits, Fisher’s *Z* transformation was applied to remove the dependency of the variance on the estimate (Equation 2), with the standard error calculated in Equation 3. The weighted mean phenotypic and genetic correlations (*r*_*w*_) were back-transformed using Equation 4,

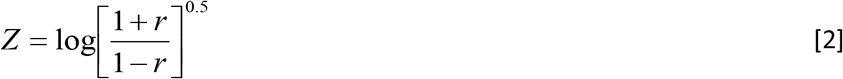

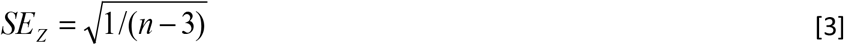

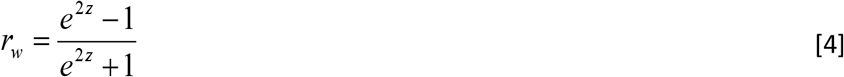

where *r* is the correlations, *n* is number of records for phenotypic correlations or number of families for genetic correlations and *z* is the weighted mean for the *Z* transformed correlations.

#### Heterogeneity

Heterogeneity among studies was assessed using three different statistical tests: *Q, H* and *I*. The *Q* statistic (Cochran 1954) tests the null hypothesis that the true parameter estimate is the same in all studies *vs*. the alternative that at least one of the estimates differs. It approximately has a Chi-square distribution with *k*-1 degrees of freedom where *k* is the number of parameter estimates. The statistic *H* different from one indicated heterogeneity of studies. In addition, the inconsistency index *I*^2^ statistic (ranging from 0 to 100%), which is defined as the percentage of observed between-study variation that is due to heterogeneity rather than chance, was also used. Significant heterogeneity was determined by *P* < 0.1 or *I*^*2*^ > 50%. Although heterogeneity was not significant for the majority of traits, random model was used to derive the weighted mean for the parameter estimate (see section on random model).

Publication bias (small study effect): To assess the effect of publication bias, the Begger’s funnel plot and Egger regression asymmetry test were used. The funnel plot visually displays the standard error of the parameter estimate of each study against its log value. The Egger test determine whether the intercept deviated significantly from zero and an asymmetric plot indicate possible publication bias, and the degree of asymmetry was tested using Egger’s test (*P* < 0.05 indicated significant publication bias)(Egger et al. 1997). Sensitivity analysis was performed by excluding studies with *z*-value greater than 3 (one study at a time) to assess the influence of the individual study on the overall results.

### 3. Estimation of weighted means of genetic parameters (heritability, common full-sib effect and correlations)

The weighted means of genetic parameters were obtained from a random effect model (Equation 5) in the metaphor package (Viechtbauer 2010). This model accounted for between- and within-study variation. Restricted maximum likelihood analysis was conducted in ASReml (Gilmour et al. 2009) to determine the relative contributions of between- and within-study variation to the weights

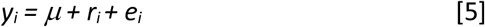

where *y*_*i*_ is the estimate of a parameter in the *i*^*th*^ study, *μ* is the mean (weighted mean of the estimates), *r*_*i*_ is the random term to account for variation between studies and *e*_*i*_ is the error term (within-study variation). The components of *r*_*i*_ and *e*_*i*_ are assumed to be normally distributed with means of zero and variances of *r*_*i*_^2^ and *V*, respectively. For heritability and common full-sib effects, *V* =*σ*_*e*_^2^/*w*_*i*_ where *w*_*i*_ = 1/(*σ*_*r*_^2^ +*i*^2^). The *V* for genetic correlations is *V* = *σ*_*e*_^2^/(*n* -3) where *n* is the number of sires. The 95% confidence interval for the estimates was calculated from *V*, with those for genetic correlations being back transformed using equation 4.

### 4. Calculation of statistical power

I considered a simplest sib design in which *N* sires are each mated to *n* unrelated dams. A further assumption was that the sire variance and error term was normally distributed. Power calculation of this design for variance component estimation followed the formula given by Lynch and Walsh (1998, page 887) as:

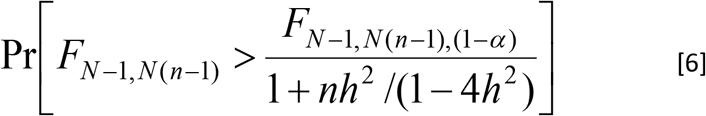

where *N* is number of sires. Each is mated with *n* dams. Alpha (*α*) is a chosen significance level (5, 1 or 0.1%) and *h*^*2*^ is the heritability for the trait in question. *F* is the test statistic having *N* degrees of freedom for the numerator and *N(n-1)* for the denominator.

I modelled four options with different numbers of sires and dams to calculate statistical power: 1) 8 sires and 16 dams, 2) 15 sires and 30 dams, 3) 30 sires and 60 dams, and 4) 45 sires and 90 dams. The ratio of males to females was based on the design of several breeding programs, although mating of one male to two females was not always successful. These parameters were used to calculate statistical power to detect a significant heritability at a probability of 95% or 99% (α = 0.05 or 0.01) (Equation 6).

## III. RESULTS

### 1. Characteristics of the data

Characteristics of the studies reviewed here are presented in Table 1. The species considered included salmonids, carps, tilapias, crustaceans and molluscs (species composition is given in Table 1). A number of recent studies have been conducted in freshwater and marine finfishes and they were classified as other fishes, including yellowtail kingfish, Atlantic charr, Asian and European seabasses, seabream, common sole, yellow perch, flounder and turbot. Five major groups of traits were examined, namely growth-related traits, carcass and flesh quality, disease resistance, reproductive performance and fitness-related traits (mainly deformity and survival).

**Table 1:** Data characteristics for reports on genetic parameters for growth related traits

| Species | No. of studies | Age of pub years relative to 2017 | Average data records | Average no of sires | Average no of dams | Design | Analytic methods |
| --- | --- | --- | --- | --- | --- | --- | --- |
| Salmonids | 40 | 11.9 | 27,933 | 104 | 69 | S, M, C | REML |
| Tilapias | 34 | 7.9 | 16,096 | 164 | 223 | S | IC, REML, $h^2_R$ |
| Carp | 8 | 8.8 | 4,208 | 166 | 279 | S, M, C | REML |
| Other fishes | 31 | 5.9 | 3,296 | 122 | 95 | S, M, C | REML |
| Crustacean | 24 | 7.0 | 41,135 | 294 | 392 | S | REML |
| Molluscs | 12 | 8.7 | 6,189 | 112 | 172 | S, M | IC, REML, $h^2_R$ |
| Across species | 150 | 8.7 | 19,357 | 199 | 264 | S, M, C | IC, REML, $h^2_R$ |

Salmonids = Atlantic salmon (*Salmo salar*), Coho salmon (*Oncorhynchus kisutch*), rainbow trout (*Oncorhynchus mykiss*), brown trout (*Salmo trutta*), Arctic char (*Salvelinus alpinus*) and European whitefish (*Coregonus lavaretus* L.)
Tilapias = Nile (*Oreochromis niloticus*), blue (*Oreochromis aureus*), red (*Oreochromis sp*) and African indigenous (*Oreochromis shiranus*) tilapia
Carps = common carp (*Cyprinus carpio*), rohu carp (*Labeo rohita*), silver carp (*Hypophthalmichthys* *molitrix*)
Other fishes include yellowtail kingfish (*Seriola lalandi*), seabream (*Sparus aurata*), Asian seabass (*Lates calcarifer*), European seabass (*Dicentrarchus labrax*), Atlantic cod (*Gadus morhua*), common sole (*Solea solea*), striped catfish (*Pangasianodon hypophthalmus*), Channel catfish (*Ictalurus* *punctatus*), turbot (*Scophthalmus maximus*), Yellow perch (*Perca flavescens*)
Crustaceans = Whiteleg shrimp (*Penaeus (Litopenaeus) vannamei*), tiger prawn (*Penaeus monodon*), giant freshwater prawn (*Macrobrachium rosenbergii*), banana prawn (*Fenneropenaeus merguensis*) and redclaw crayfish (*Cherax quadricarinatus*)
Molluscs = Pacific oyster (*Crassostrea gigas*), Sydney rock oyster (*Saccostrea glomerata*), tropical oyster (*Saccostrea cucullata*), Pearl oyster (*Pinctada fucata*), silver-lip Pearl oyster (*Pinctada* *maxima*), Chinese Pearl oyster (*Pinctada martensii*), Chilean mussel (*Mytilus chilensis*), Mediterranean mussel (*Mytilus galloprovincialis*), small abalone (*Haliotis diversicolor*), South African abalone (*Haliotis midae*), red abalone (*Haliotis rufescens*), tropical abalone (*Haliotis asinina*), scallop (*Placopecten magellanicus*), Japanese scallop (*Patinopekten yessoensis*), catarina scallop (*Argopecten* *ventricosus*) and Manila clam (*Ruditapes philippinarum*)
S = Single pair mating with known parental identities to maintain pedigree using conventional physical tags (Passive Integrated Transponder, PIT tags for fish and visible implant elastomer, VIE for crustaceans), M = Mass spawning using DNA markers for parentage assignment, and C = Combined conventional and molecular genetic pedigree methods
IC = Intraclass correlation, REML = Restricted Maximum Likelihood Method under the mixed model framework, and $h^2_R$ = realised heritability

The average sample size (number of individuals) of the studies included in this review ranged from 3,296 to 27,933 (Table 1). The average number of sires and dams (or number of families) across populations were 199 and 264, respectively. A majority of the studies applied single-pair mating (with a hierarchical nested mating design with a ratio of one male to 2 females) or incomplete (rectangular) factorial mating by set (e.g. male 1 mated with females 1 and 2, male 2 with females 2 and 3, and so on to male *n* with female 1 and *n*). Mass spawning also was practised for some species, mainly in new aquaculture species including yellowtail kingfish, seabream, abalone and blue mussel, where microsatellite DNA markers were used for parentage assignment. There is an increasing trend in the number of studies that have used DNA markers for parentage assignment and pedigree construction. Conventionally, individual identification in fish involved use of Passive Integrated Transponder (PIT) tags, whereas visible implant elastomer (VIE) (or visible implant alphanumeric) in combination with eye tags were used in crustacean species. A wide array of testing environments during grow-out were used, predominantly ponds or cages. Only three studies employed recirculating or intensive culture systems.

With the pedigree information available in almost all studies (96.9%), Restricted Maximum Likelihood (REML) method applied to a single or multiple trait mixed model was performed to estimate genetic parameters. Some studies reported realised heritability from selection experiments (1.5%). A small proportion of the *h*^*2*^ estimates was also obtained from conventional intraclass correlation analysis (only three studies).

### 2. Summary statistics

Body weight is frequently recorded in selective breeding programs for aquaculture species (*n* =193). There is a growing interest regarding genetic improvement for disease resistance (*n* =55). Only a limited number of studies investigated flesh/eating quality attributes (*n* =18), reproduction traits (*n*=9) and morphological deformity (*n*=11) in aquaculture species. Limited samples sizes were observed for quality, reproductive and fitness-traits, mainly because they are expensive or difficult to measure. The coefficients of variation for reproductive traits were substantially larger than growth and carcass/flesh quality characteristics (results not tabulated). The mean and standard deviations are trait-, population-, or environment-specific because market weights of aquaculture species are different or harvesting time to make measurements varied greatly among species; hence, basic statistics for traits studied were not summarised.

### 3. Factors affecting genetic parameters

The General linear model (GLM procedure) to examine the significance of fixed effects [including species (salmonids, carps, tilapias, crustaceans, molluscs and other fishes), experimental design (single-pair mating vs. mass spawning), statistical analytic method (REML, realised heritability and intraclass correlation analysis) and age of the publication (or assemble data collected) relative to 2017] showed that only experimental design and analytic method were statistically significant for body weight (*P* < 0.01 and 0.05, respectively) (Table 4).

### 4. Heritability

The number of studies, weighted mean heritabilities (± standard errors, S.E), between-study variances, *Q*-statistics and 95% confidence intervals (CI) of the estimate for different traits grouped for all species are presented in Tables and 2. The results with specificities for some fish species are considered below.

**Table 2:** Number of literature estimates (*N*), weighted mean heritability (*h*^*2*^), between study variance 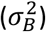, Q statistics and 95% confidence interval (CI) of the heritability estimate for economic traits across aquaculture species.

| Traits | $N$ | $h^2 \pm s.e.$ | $\sigma_B^2$ variance | $Q$-statistic | 95% CI of $h^2$ |
| --- | --- | --- | --- | --- | --- |
| **Growth traits** |  |  |  |  |  |
| Body weight | 193 | 0.30±0.02*** | 0.14×10<sup>-5</sup> | 82.0 | 0.27 – 0.33 |
| Feed utilisation | 14 | 0.23±0.10* | 0.92×10<sup>-1</sup> | 5.5 | 0.04 – 0.42 |
| **Carcass and flesh quality traits** |  |  |  |  |  |
| Fillet weight | 8 | 0.33±0.10*** | 0.10×10<sup>-6</sup> | 0.69 | 0.14 – 0.51 |
| Fillet yield | 3 | 0.20±0.13<sup>ns</sup> | 0.40×10<sup>-1</sup> | 1.31 | -0.05 – 0.45 |
| pH | 1 | 0.15±0.14<sup>ns</sup> | 0.01×10<sup>-6</sup> | 0.00 | -0.13 – 0.32 |
| Colour | 11 | 0.18±0.07* | 0.51×10<sup>-1</sup> | 1.84 | 0.04 – 0.32 |
| Protein content | 5 | 0.05±0.14<sup>ns</sup> | 0.63×10<sup>-3</sup> | 0.01 | -0.22 – 0.33 |
| Fat content | 18 | 0.17±0.07* | 0.10×10<sup>-6</sup> | 3.98 | 0.03 – 0.31 |
| **Reproductive traits** |  |  |  |  |  |
| Sexual maturity | 9 | 0.21±0.09* | 0.10×10<sup>-6</sup> | 1.1 | 0.04 – 0.38 |
| Female weight | 7 | 0.35±0.13** | 0.49×10<sup>-2</sup> | 1.66 | 0.11 – 0.60 |
| Fecundity | 9 | 0.14±0.12<sup>ns</sup> | 0.18×10<sup>-6</sup> | 0.03 | -0.10 – 0.37 |
| Fry weight | 2 | 0.09±0.13<sup>ns</sup> | 0.19×10<sup>-4</sup> | 0.02 | -0.16 – 0.35 |
| Fry survival | 6 | 0.08±0.14<sup>ns</sup> | 0.89×10<sup>-4</sup> | 0.16 | -0.19 – 0.36 |
| **Disease resistance** |  |  |  |  |  |
| Virus | 27 | 0.27±0.04*** | 0.58×10<sup>-6</sup> | 11.9 | 0.18 – 0.35 |
| Bacteria | 21 | 0.32±0.05*** | 0.18×10<sup>-6</sup> | 13.5 | 0.23 – 0.41 |
| Parasites | 15 | 0.20±0.06** | 0.16×10<sup>-6</sup> | 4.5 | 0.09 – 0.31 |
| Others | 5 | 0.20±0.11<sup>ns</sup> | 0.68×10<sup>-2</sup> | 2.3 | -0.01 – 0.42 |
| All diseases | 55 | 0.26±0.02** | 0.20×10<sup>-5</sup> | 67.5 | 0.22 – 0.30 |
| **Fitness related traits** |  |  |  |  |  |
| Deformity | 11 | 0.18±0.11<sup>ns</sup> | 0.43×10<sup>-3</sup> | 0.84 | -0.04 – 0.39 |
| Survival | 31 | 0.14±0.03*** | 0.45×10<sup>-5</sup> | 21.0 | 0.08 – 0.21 |

\* $P < 0.05$ , \*\* $P < 0.01$ , \*\*\* $P < 0.001$ and ns = non-significance

#### a. Growth related traits

Regardless of whether analyses were conducted separately for each species or for all species together, the weighted heritabilities for growth-related traits (body weight) were moderate (0.22 – 0.33), suggesting that selective breeding is likely to prove successful for growth characters. A number of studies calculated body shape based on body measurements (weight/ length or depth/length) (Trọng et al. 2013a) or from image analysis (Blonk et al. 2010). The average heritability estimate for body shape was low to moderate, suggesting that changing this trait in aquaculture species is possible as predicted from selection index theory (Nguyen et al. 2007). Prchal et al (2018) also informed that selection for a trait connected to a trait of interest may have additional effect on body shape. In addition to growth-related traits, the heritability for food utilisation efficiency (FUE) was moderate (0.23 ± 0.10, 95% CI = 0.04 to 0.42). Direct selection for ratio traits (i.e., FUE) is, however, difficult. This trait can be sometimes improved indirectly through selection for increased growth or reduced body fat (Verdal et al. 2017).

#### b. Carcass and flesh/eating quality

A carcass trait of paramount importance in fish is fillet yield or edible tail meat for shrimp (Hung and Nguyen 2014). In both fish and shrimp, fillet or edible meat weight is moderately heritable (*h*^*2*^ = 0.33 ± 0.10), whereas the heritability estimate for fillet yield (or percentage of fillet over body weight) was of lower magnitude (mean *h*^*2*^ = 0.20 ± 0.13, P > 0.05). Flesh and eating quality are complex characteristics and they are strongly influenced by environmental factors before slaughter, such as fasting, handling and post-mortem processing techniques (Hultmann et al. 2016). This is indicated by the low and insignificant heritability for the important flesh and eating quality attributes (pH, protein content) included in this review (Table 2). Almost all studies engaged total fat content of fillet (Hamzah et al. 2014b); information on fat deposits in different body compartments/locations could have genetically different patterns (Kause et al. 2002; Tobin et al. 2006). Across the measures of fat content reported in the literature, this trait had moderate heritability (mean *h*^*2*^ = 0.17, *P* < 0.05). Flesh colour in some fishes is an attribute of biological importance because it is positively correlated with pH, water holding capacity and eating characteristics (mean *h*^*2*^ = 0.18, P < 0.05) and also because it affects market value. For example, deeply red salmon fillets are more highly valued than whitish ones. In fish, flesh colour was subjectively measured based on a scoring scale or objectively measured using a colour wheel or instrument. Norris and Cunningham (2007) reported higher heritability for objective than subjective measurements of flesh colour in Atlantic salmon. Based on a binary scale (visual observation of red or light colour), Nguyen *et al*. (2014) suggested that there is a possibility for improving red body colour of both raw and cooked banana shrimp.

Other flesh and eating quality characteristics reported in the literature exhibited additive genetic variation in muscle fibre types or fibre density in rainbow trout, with the estimate of heritability ranging from 0.10 to 0.33 (Vieira et al. 2007). Two studies reported heritability for fatty acid composition in Nile tilapia (Nguyen et al. 2010b) and Atlantic salmon (Leaver et al. 2011) and another in Whiteleg shrimp (Nolasco-Alzaga et al. 2018). For these flesh/eating quality attributes, the number of studies are very limited (<3); hence, the weighted heritability was not calculated.

#### c. Disease resistance

Heritability estimates for disease resistance were mainly reported from challenge test studies for three types of pathogens (parasites, bacteria and virus). In practice, the resistance was measured as survival rate of the challenged animals with *LD*_*50*_ to a particular time point, after a period of 45-50 days. Only one study proposed an alternative approach to estimate disease resistance, using qPCR analysis to measure HPV viral load of individual banana shrimp (*Fenneropenaeus merguiensis*)(Knibb et al. 2015). I calculated weighted mean heritability separately for each disease type (bacterial, parasitic and viral) and across all the three types of pathogens (Table 2). The results showed that a significant heritable genetic component exists for a variety of diseases in fishes, shrimps and molluscs. The magnitude of the heritability estimates was not significantly different among the three disease types (Tables 2). Three studies (Camara et al. 2017; Degremont et al. 2015; Liang et al. 2017) estimated heritability for disease resistance in mollusc (2 in oysters and 1 in clam).

Six studies reported the heritability for indicators of stress resistance (i.e. cortisol level) or hypoxia and oxidative stress resistance. The weighted heritability for these indicators of stress tolerance was moderate (0.20 ± 0.11) but not significant (*P* > 0.05).

Across species, the moderate level of heritability for resistance against a range of diseases and environmental stressors suggests that these traits would respond effectively to selection. Improving disease resistance is necessary to reduce mass mortality and associated loss of weight and economic opportunities for the aquaculture sector worldwide. Novel approaches (e.g., markers-assisted or genome-based selection) should be considered as conventional method based on challenge tests is associated with ethic issues or due to some notifiable status of some diseases or other veterinarian measures (Trinh et al. 2019).

#### d. Reproduction

Reproductive traits reviewed here included sexual maturity, female weight prior to spawning, egg size, fecundity (egg numbers or egg volume), larval survival and fry weight. In selective breeding programs for many aquatic species, status of sexual maturity during the grow-out period was recorded as a binary character and analysed using both linear and threshold mixed models. Across the studies and species examined here, sexual maturity exhibited a low to moderate heritability (weighted mean *h*^*2*^ of 0.21 and confidence interval ranging from 0.04 to 0.38). However, the heritability for age at sexual maturity was high (Crandell and Gall 1993; Gjerde 1984). The mean heritability for female weight was moderate (*h*^*2*^ = 0.35). By contrast, the heritabilities estimated for other reproductive traits were low and non-significant (Table 2). There were no differences in the heritability estimates for reproductive characteristics between fishes (Gima et al. 2014; Thoa et al. 2017; Trọng et al. 2013c) and shrimps (Caballero-Zamora et al. 2015; Macbeth et al. 2007). The consistently low heritability for reproductive and fitness-related traits across aquatic species suggests that response to selective breeding to improve these characters, albeit possible, may be slow and difficult to achieve. In addition to the five important reproductive traits reviewed here, low to moderate heritability was reported for gonad weight/gonad somatic index (Charo-Karisa et al. 2007) or spawning interval in Nile tilapia (Trọng et al. 2013b). The heritability for egg diameter or hatching rate was low (essentially not different from zero). Generally, reproductive traits are difficult to be investigated for heritability, as these characters are greatly affected by environmental and other variables (e.g., health status of fish, level of management of artificial reproduction) that bias the genetic variation estimates. Thus this area of research deserves further studies across aquaculture species.

#### e. Fitness-related traits

Two fitness related traits studied here were survival and deformity. Survival during grow-out showed a low but significant additive genetic variation, with the heritability mean estimate of 0.14 (95% confidence interval, CI from 0.08 to 0.21, *P* < 0.001) across species. Existence of the heritable (additive) genetic component for survival during the early phase of growth (i.e., before physical tagging or 1-2 months after hatching) has been reported for tilapia (Thoa et al. 2015) and rainbow trout (Vehviläinen et al. 2012).

There are various forms of deformity in fishes such as lower jaw protrusion, nasal erosion, opercula distortion, cataracts, spinal abnormalities (lordosis, kyphosis, scoliosis, and ankylosis) as well as mouth and fin malformations. Estimates of heritability were reported for a range of deformity measures or for a specific type of deformity, such as humpback anterior dorsal fin, humpback posterior dorsal fin and shortened tail fin in Atlantic salmon (Glover et al. 2005) or lordosis, cyphosis, scoliosis and vertebral fusion in European seabass (Karahan et al. 2013). Synthesised results from the literature showed that different measures of deformity can be considered as separate traits, including studies in rainbow trout (Vehviläinen et al. 2012) and kingfish (Nguyen et al. 2016). However, a range of measures of deformity was pooled in this study, mainly due to the limited sample size (small number of studies). The weighted heritability estimate for this trait was low and not significant (weighted mean *h*^*2*^ = 0.18 ± 0.11, *P* > 0.05).

#### f. New traits

Novel traits considered in this review included behaviour, socially indirect genetic effects and uniformity. A detailed description of measurement methods is given in Supplementary N.4. Recent studies have attempted to qualify the level of genetic variation for these new traits, with the heritability ranging from 2 to 10% for uniformity (Sae-Lim et al. 2015) and direct or indirect social genetic traits (Nielsen et al. 2014). There is also evidence of genetic variability in personality-related characters in rainbow trout (Millot et al. 2014). However, the heritability estimates for these behavioural traits ranged from low in brown trout (Kortet et al. 2014) to moderate in Atlantic cod (Drangsholt et al. 2014). A high heritability for boldness (risk-taking score) was, however, reported for European seabass (Ferrari et al. 2016). There is a need for further studies to support a reliable systematic meta-analysis for novel traits across farmed aquaculture species of economic importance.

### 5. Common full-sib effects

In aquatic species, different experimental designs have been utilized to estimate heritability-related parameters for a range of traits of economic importance. Common full-sib effects (*c*^*2*^) were a result of separate family rearing of about one to two months until fish/shrimps reach a suitable size for physical tagging (5-10 g in fish and 1-2 g in shrimps). The *c*^*2*^ includes both common environmental and maternal effects, accounting for approximately 5 – 14% of the total phenotypic variance across species (Table 3). The *c*^*2*^ effects show a tendency to diminish in later phase of growth. For carcass and flesh quality traits, these effects were generally not significant. The *c*^*2*^ effects were not significant for eating quality characteristics or fillet fatty acid composition. Only one study (Hamzah et al. 2014a) reported *c*^*2*^ effects for reproductive traits, but the estimates were essentially zero. Maternal genetic effects have not been reported for disease resistance or measures of deformity.

**Table 3:** Number of literature estimates (*N*), weighted mean maternal and common environmental effects (*c*^*2*^), between study variance 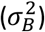, *Q* statistics and 95% confidence interval (CI) for economic traits across aquaculture species.

| Traits | $N$ | $c^2 \pm s.e.$ | $\sigma_B^2$ variance | $Q$-statistic | 95% CI of $c^2$ |
| --- | --- | --- | --- | --- | --- |
| **Growth traits** |  |  |  |  |  |
| Body weight | 45 | 0.10±0.02** | 0.97×10<sup>-5</sup> | 12.7 | 0.05 – 0.14 |
| Feed utilisation | 2 | 0.14±0.17<sup>ns</sup> | 0.15×10<sup>-6</sup> | 0.01 | -0.18 – 0.47 |
| **Carcass and flesh quality traits** |  |  |  |  |  |
| Fillet weight | 2 | 0.07±0.09<sup>ns</sup> | 0.36×10<sup>-2</sup> | 0.15 | -0.10 – 0.25 |
| **Reproductive traits** |  |  |  |  |  |
| Sexual maturity | 1 | 0.090±0.04<sup>ns</sup> | 0.00×10<sup>-5</sup> | 0.01 | -0.18 – 0.26 |
| **Fitness related traits** |  |  |  |  |  |
| Survival | 6 | 0.04±0.05<sup>ns</sup> | 0.10×10<sup>-6</sup> | 0.32 | -0.05 – 0.13 |

\*\* $P < 0.01$ and ns = non-significant

**Table 4:** *F*-value of factors included in the analysis of heritability for growth related traits studied

| Traits | *N* | *R*<sup>2</sup> | Species | Design | Analysis<br>method | Assemble<br>data<br>colected |
| --- | --- | --- | --- | --- | --- | --- |
| Body weight | 193 | 0.65 | 1.4 | 3.6** | 3.1* | 0.3 |
| Survival | 31 | 0.93 | 2.4 | 0.2 | 0.5 | 0.1 |
| Sexual maturity | 9 | 0.87 | 3.9 | n.a. | n.a. | 6.6 |

\**P*<0.05, \*\**P*<0.01.
These effects on other traits (feed utilisation, flesh quality, reproduction, disease resistance and deformity) were also not significant

### 6. Heritabilities in contrasting environments

Because expression of quantitative traits depends in part upon environmental factors, heritabilities estimated in diverse environments may differ. Figure 2 presents standardized mean differences in heritabilities for body weight between contrasting environments. There was non-significant difference in the magnitude of heritability between the environments, suggesting that there is no (or little) reduction in the additive genetic variance of body weight in the production environment compared with the nucleus. Additionally, previous studies (Nguyen 2016; Sae-Lim et al. 2016) showed that the between-environment genetic correlations for homologous body traits was positive and high when the environments used in the nucleus and production were similar. However, when the two environments differed, the genetic correlation estimates was low or moderate, suggesting that the G×E interaction effects are potentially important for complex traits in aquaculture species.

**Figure 1:**
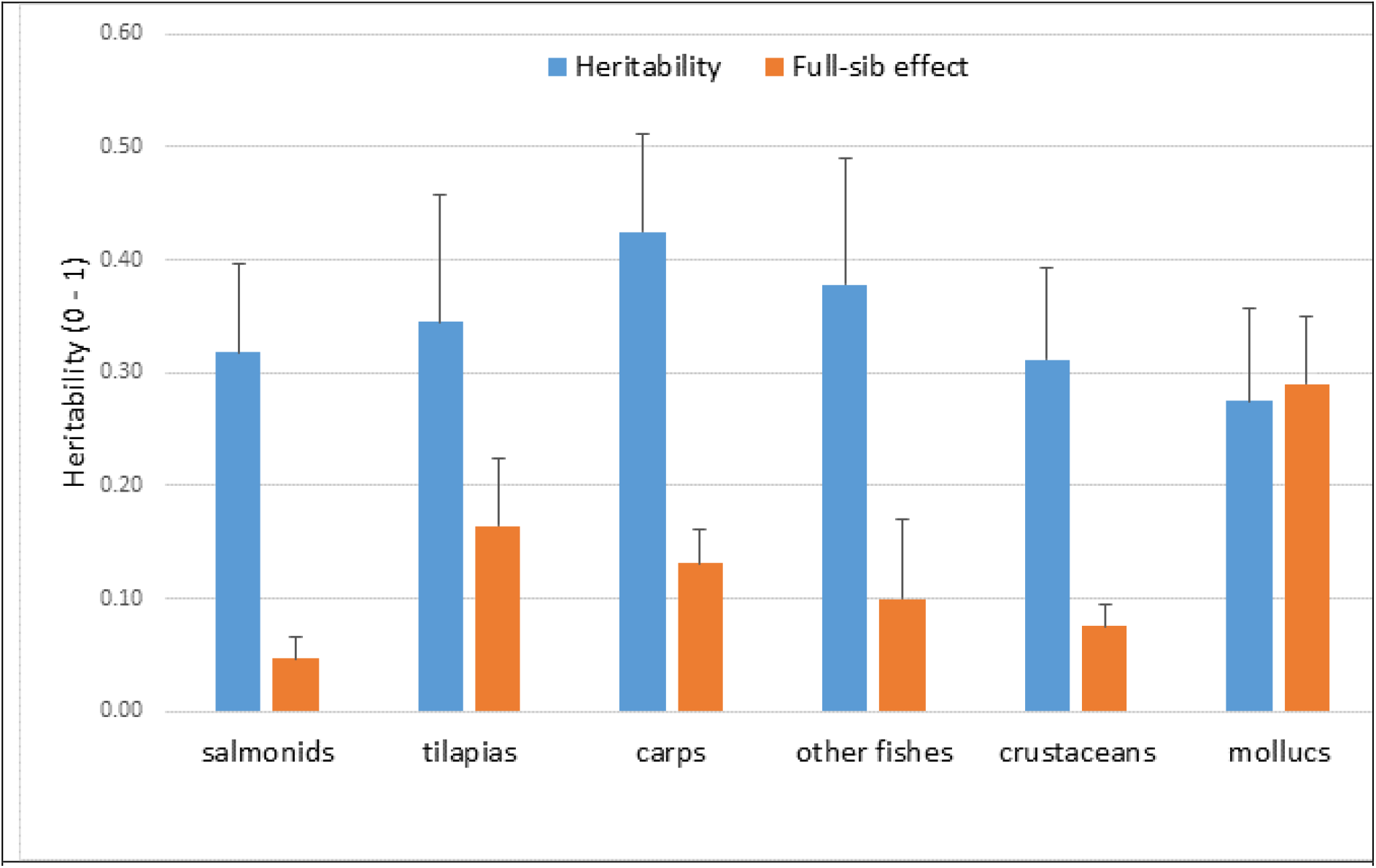
Weighted mean heritability (*h*^*2*^) for body weight and the maternal and common environmental full-sib effects (*c*^*2*^) in six groups of aquaculture species studied.

**Figure 2:**
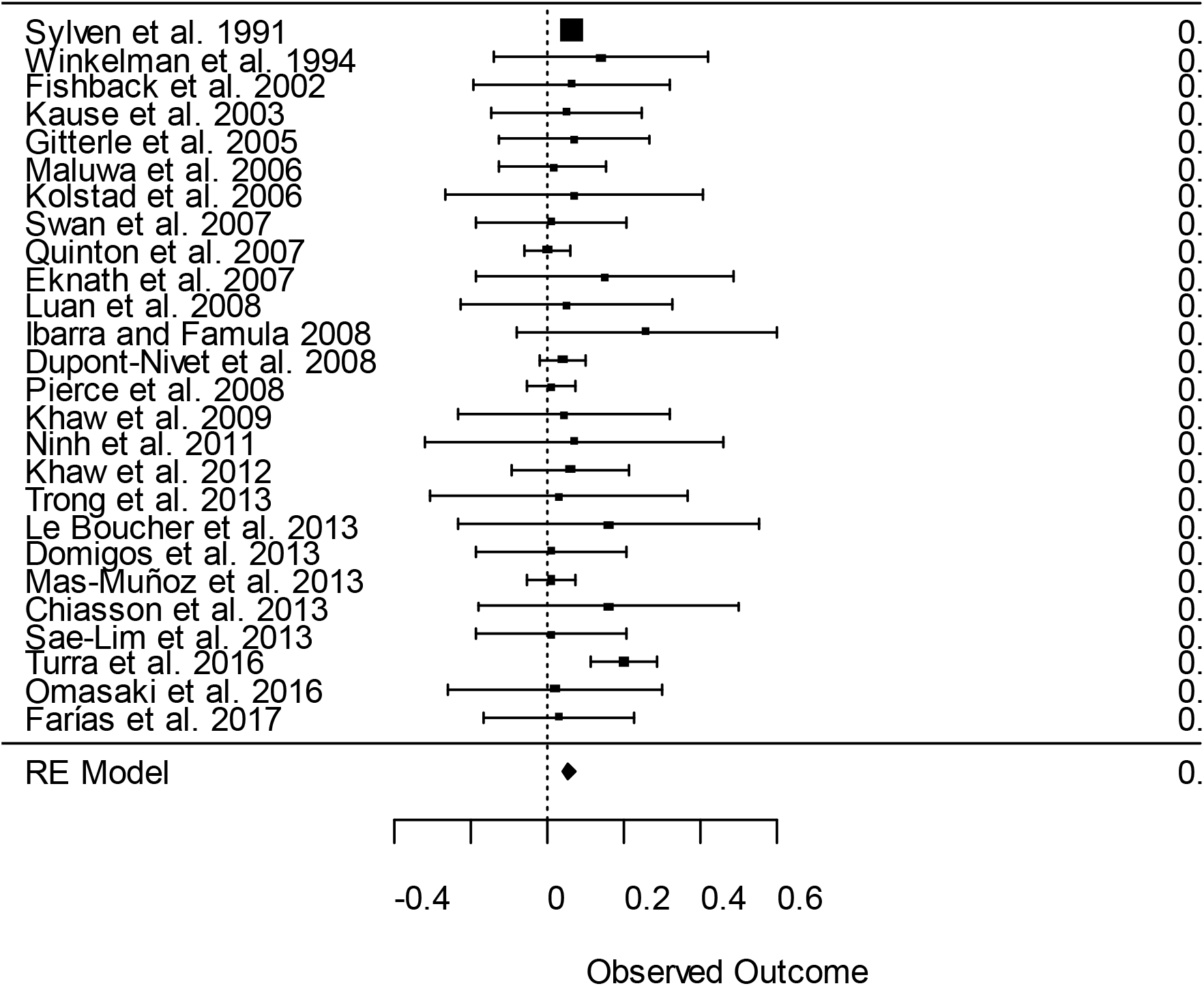
Forest plot of heritability difference with 95% confidence interval between contrasting environments (*Q*-statistic = 22.530, *p*-value = 0.605) and between-study variance = 0.03. RE = Random effect model. Solid square = weighted mean

### 7. Genetic correlations

#### a. Correlations between body and carcass/flesh quality

Certain traits of breeding interest are correlated with one another. Knowledge of such correlations is needed to design purposeful breeding programs. Genetic correlations (*r*_*g*_) between flesh quality and body weight (the sole selection criterion of many breeding programs) are shown in Table 5. The genetic correlations between fillet fat content and body weight were moderate to high and mostly positive (mean *r*_*g*_ = 0.51 ± 0.06), except for the negative estimate reported in rainbow trout by Kause et al. (2002). The estimated correlation between moisture and body weight was positive in salmonids and tilapias, but negative in rainbow trout. Flesh or body colour exhibited a moderate to high positive correlation with body weight in Atlantic salmon (Quinton et al. 2005) and banana shrimp (Nguyen et al. 2014).

**Table 5:** Number of literature estimates (*N*), weighted mean genetic correlations of flesh quality traits with body weight (***r***_***g***_), between study variance 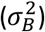, *Q* statistics and 95% confidence interval (CI) of the genetic correlation estimate

| Traits | $N$ | $r_g$ | $\sigma_B^2$ variance | $Q$-statistic | 95% CI |
| --- | --- | --- | --- | --- | --- |
| Protein | 2 | 0.02 | 0.02 | 13 | -0.17 – 0.21 |
| Fat | 12 | 0.51*** | 0.16 | 2474 | 0.33 – 0.67 |
| Moisture | 4 | -0.19 | 0.02 | 81 | -0.32 – -0.06 |
| pH | 1 | 0.01 | n.a. | n.a. | n.a. |
| Colour | 6 | 0.31** | 0.29 | 2605 | -0.08 – 0.62 |

\*\*P < 0.01 and \*\*\*P < 0.001

#### b. Correlations between body weight and disease resistance

Genetic correlations between body weight and disease resistance were either positive or non-significant in the literature (Table 6). Some studies, however, also reported negative (i.e. antagonistic) genetic association between growth and disease resistance, for example ranging from - 0.01 to -0.33 in rainbow trout (Henryon et al. 2002) or -0.54 to -0.66 in Pacific white leg shrimp (Gitterle et al. 2005). Recent studies showed weak or non-significant genetic relationship between the two traits in rainbow trout (Yáñez et al. 2014) or shrimp (Phuthaworn et al. 2016). The weighted mean genetic correlation between the two traits (weight and disease resistance) was not significant (Table 6). When the analysis was conducted separately for each type of pathogen, a significant genetic correlation was observed between body weight and resistance to parasites (*r*_*g*_ = 0.23).

**Table 6:** Number of literature estimates (N), weighted mean genetic correlations of disease resistance with body weight (***r***_***g***_), between study variance 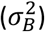, *p*-value of *Q* statistics and 95% confidence interval (CI)

| Traits | $N$ | $r_g$ | $\sigma_B^2$ variance | $Q$-statistic | 95% CI |
| --- | --- | --- | --- | --- | --- |
| Bacteria | 8 | 0.09 | 0.04 | <0.001 | -0.03 – 0.21 |
| Virus | 8 | 0.02 | 1.60 | <0.001 | -0.63 – 0.68 |
| Parasites | 10 | 0.24* | 0.19 | <0.001 | -0.12 – 0.59 |
| Across types of pathogens | 26 | 0.13 | 0.72 | <0.001 | -0.07 – 0.33 |

\*P < 0.05

#### c. Correlations between body weight and deformity

The reported estimates of genetic correlations between deformity and growth traits varied from negative (Gjerde et al. 2005) to positive (Karahan et al. 2013) or non-significant (Kolstad et al. 2006)(Figure 3). Across the studies, the estimate was not significant (*r*_*g*_ = -0.14, CI =-0.56 to 0.29, *P* > 0.05). However, the genetic relationship between body weight and deformity varied with growth phase and trait definitions, e.g. fast-growing rainbow trout fingerlings are prone to skeletal deformities, while body weight and cataracts in the adult stage were negatively correlated (−0.52 to - 0.62)(Vehviläinen et al. 2012). Nguyen et al. (2016) report that the degree of co-inheritance of deformity varied with trait combinations (i.e. positive between weight and jaw malformations but negative between weight and operculum). There is still very limited information regarding the genetic correlation of deformity with traits of economic importance, suggesting that a detailed recording of deformity characteristics would be necessary to characterize the genetic architecture of malformation in aquaculture populations reared in different culture environments.

**Figure 3:**
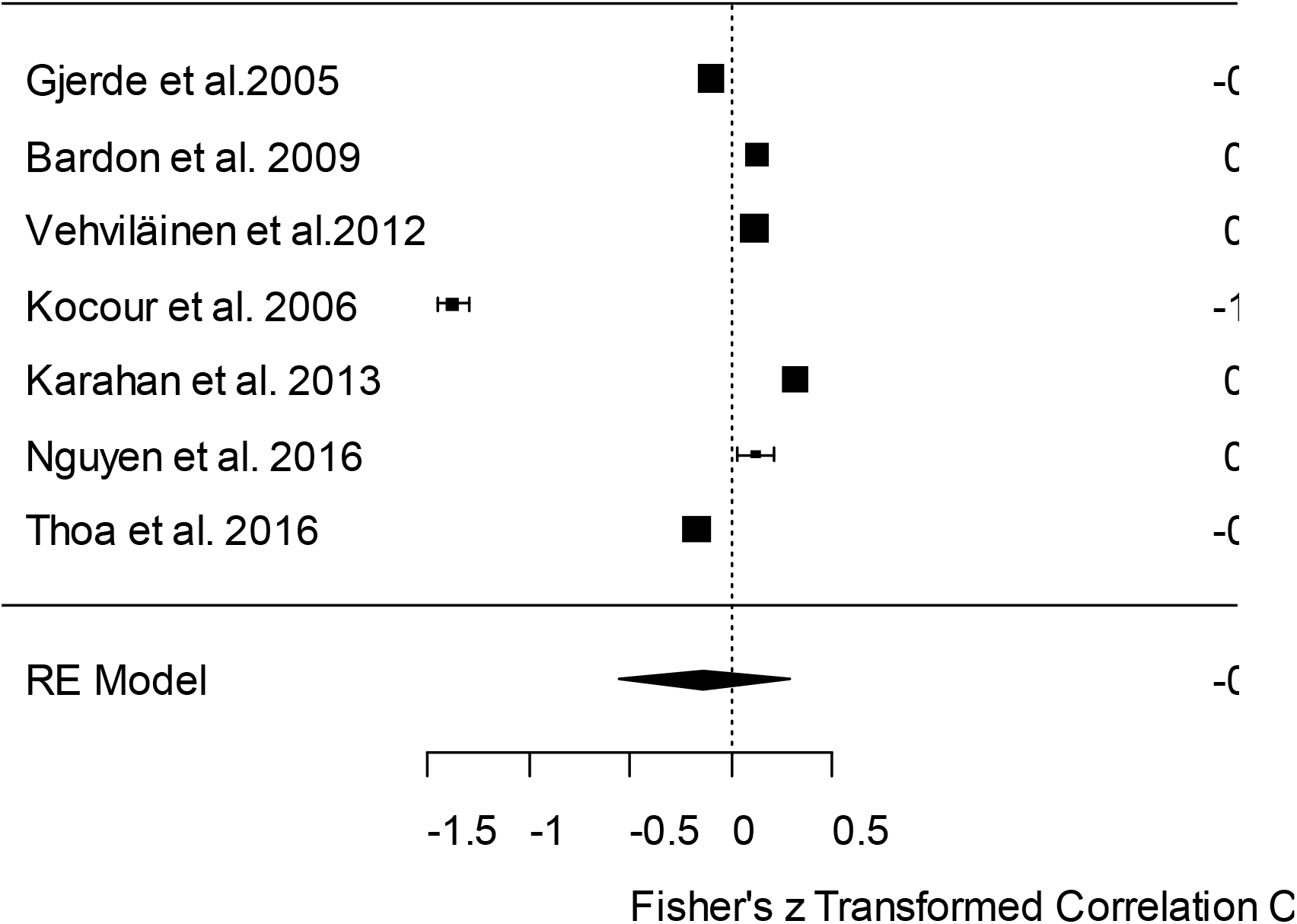
Weighted genetic correlations (r_g_ with 95% confidence interval) of body weight with measures of deformity in fish and shrimp (between-study variance = 0.32 and P > 0.05). RE = Random effect model

#### d. Correlation between weight and reproduction

The genetic correlation of body weight during grow-out and fecundity-related traits in females was moderate and positive (Table 7). Selection for high growth is thus expected to bring about favourable changes in reproductive performance (fecundity and fry weight) of the aquatic animals, thereby, leading to greater production of fry and fingerlings for marketing. This is important only in species or culture systems where the egg is the final product (e.g., sturgeon or salmon caviar). However, interpretation of these results should be with caution because the relationship between body weight and relative fecundity was not available in the literature. Further, the genetic relationship between body weight and number of fry at hatching or fry mortality during the early phase of rearing was weak and not significant in Coho salmon (Gall and Neira 2004), Nile tilapia (Hamzah et al. 2014a) or giant freshwater prawn (Vu and Nguyen 2019). In summary, genetic relationships between reproductive traits and growth deserve further studies, due to the limited number of studies available in the literature as well many other factors involved, e.g., age of maturation.

**Table 7:** Number of literature estimates (*N*), weighted mean genetic correlations of reproductive traits with body weight (***r***_***g***_), between study variance 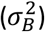, *Q* statistics and 95% confidence interval (CI)

| Traits | $N$ | $r_g$ | $\sigma_B^2$ variance | $Q$-statistic | 95% CI |
| --- | --- | --- | --- | --- | --- |
| WBS | 3 | 0.31*** | 0.00 | 0.44 | 0.29 – 0.33 |
| ES | 3 | 0.36** | 0.05 | 213 | 0.12 – 0.56 |
| TNE | 5 | 0.20* | 0.04 | 212 | 0.02 – 0.36 |
| EV | 2 | 0.50** | 0.14 | 389 | 0.04 – 0.79 |
| NFH | 3 | 0.04<sup>ns</sup> | 0.01 | 27 | -0.04 – 0.18 |

WBS= weight before spawning, ES= egg size (measured as diameter or volume), TNE= total number of eggs, EV= total volume of eggs, NFH= number of fry hatched
\* $P < 0.05$ , \*\* $P < 0.01$ , \*\*\* $P < 0.001$ and ns = non-significant

#### e. Correlation between weight and new traits

Currently, genetic associations of growth (or body weight) with behavioural and functional traits are not well documented. This is an important area that merits future studies to support the possibility of multi-trait selection and to gain knowledge to control unexpected changes from selection programs for high productivity.

### 8. Sample size

A parameter-estimation experiment needs sufficient power in order to yield a reliable estimate for design of an effective breeding program. The sample sizes required to obtain sufficient statistical power to detect significant heritability are given in Table 8. The statistical power depends on the level of heritability (low, moderate or high) and the probability of detecting significant heritabilities (95 or 99%). Statistical power increases with the number of sires and dams in the pedigree. For a given sample size, statistical power is higher for greater heritabilities. Statistical power also differs with levels of probability associated with detecting significant heritability. Across the three scenarios studied, with the hierarchical nested design a minimum 30 sires and 60 dams are required to obtain sufficient (>80%) power to detect a significant heritability estimate. Fewer than 30 families are not recommended because under this mating (nested) scheme, the likelihood of detecting significant heritability was low, ranging from 3.6 to 34.6%.

**Table 8:** Number of sires and dams required in a nested design (one male is mated two females)

|  | Option 1 |  |  |  | Option 2 |  |  |  | Option 3 |  |  |  |
| --- | --- | --- | --- | --- | --- | --- | --- | --- | --- | --- | --- | --- |
|  | *P* = 0.05 |  | *P* = 0.01 |  | *P* = 0.05 |  | *P* = 0.01 |  | *P* = 0.05 |  | *P* = 0.01 |  |
| Sires | 15 | 15 | 15 | 15 | 30 | 30 | 30 | 30 | 45 | 45 | 45 | 45 |
| Dams | 30 | 30 | 30 | 30 | 60 | 60 | 60 | 60 | 90 | 90 | 90 | 90 |
| *h*<sup>2</sup> | 0.07 | 0.20 | 0.07 | 0.20 | 0.07 | 0.20 | 0.07 | 0.20 | 0.07 | 0.20 | 0.07 | 0.20 |
| **Power** | **35.7** | **82.8** | **16.5** | **67.1** | **87.6** | **99.8** | **73.4** | **99.8** | **99.6** | **100** | **98.4** | **99.9** |

## IV. DISCUSSION

This is the first systematic meta-analysis of genetic parameters for aquatic animal species. The heritabilities, common full-sib effects and correlations provide information advancing our understanding of genetic architecture of economically important traits in aquatic animal species. My results also help to fill a gap in our knowledge regarding inheritance and relationships among quantitative traits, providing knowledge to address challenges regarding genetic improvement of aquaculture species. In addition, this systematic meta-analysis identified limitations and proposed suggestions for future studies as well as possibilities to utilise advanced statistical methods and genome sequence technologies to understand complex quantitative traits of aquatic species. Specifically, the results synthesised from this study provide answers to the following questions:

### 1. Does the quantitative genetic basis of complex traits differ?

The weighted mean of heritability estimated from the meta-analysis (Figure 1 and Table 2) showed that quantitative genetic architecture differed among the five groups of traits reviewed. Growth- and carcass-related characters had moderate to high heritabilities. On the other hand, traits related to quality (fat content, colour) or fitness (reproduction: fecundity, early survival) are lowly or moderately heritable. Disease resistance to different types pf pathogens (bacterial, viral and parasitic) exhibited substantial heritable genetic variance in a range of aquaculture species (Ødegård et al. 2011). My findings are consistent with the expectation of quantitative genetic and evolutionary theory as well as with observations in terrestrial farmed animals and model species (Safari et al. 2005).

### 2. Can the traits studied show response to selection?

With abundant genetic variation in traits related to growth, genetic improvement of these characters should prove successful, as demonstrated in several selection programs across aquatic species ranging from fishes (Hamzah et al. 2014c) to crustaceans (Hung et al. 2013b) and molluscs (Liu et al. 2015). Across aquaculture species, genetic gain has ranged from 8 – 22% per generation (Gjedrem and Rye 2016; Nguyen 2016). On the other hand, selective breeding for other traits, such as reproduction, fitness and deformity, may be more difficult; hence, systematic data recording of sibling information from multi-generational, in-depth pedigree populations is needed to obtain reliable estimated breeding values for selection candidates. Genetic improvement for traits that require sacrifice of animals such as flesh/eating quality or disease resistance is still difficult. Further, only sibling information can be used, thereby reducing accuracy of estimated breeding values and selection response. In these cases, genomic selection can potentially speed up genetic progress in traits that are difficult or expensive to measure (Boison et al. 2019).

### 3. Are there heritable additive genetic components for new traits?

In addition to the existence of useful additive genetic variance for fitness characters studied here (maturity, survival, reproduction), the proportion of genetic relative to total phenotypic variance for behavioural, social and immunological traits ranges from 0.03 to 0.45 (Ferrari et al. 2016; Kortet et al. 2014). These traits were not included in the meta-analysis, as only a limited number of studies are available in the literature and, the data mostly were recorded from a limited number of animals in shallow pedigrees of only one or two generations. Large-scale measurements for behavioural/functional and adaptive traits (e.g. stress tolerance to environmental factors) are still challenging due to the costs involved and sophistication of data recording, using video or laboratory-based tests. I also attempted to calculate the weighted heritability for these traits, but the estimates had large standard errors and thus they were not significantly different from zero. To understand quantitative genetic basis of traits that are expensive and difficult to measure (e.g., flesh and eating quality or disease resistance) in aquaculture species, development of new measurement methods is required to enable routine large-scale data recording for species of commercial importance (see section 4.7).

### 4. Does the heritability of traits differ between environments?

Heritability reported in various culture environments is mainly for growth related traits (weight or daily gain). Although magnitude of the heritability estimate varied with environments, the weighted effect size was small and not significant (Figure 2). However, there are exceptions in the literature where there is a reduction in the heritability estimates for growth traits in poorly managed culture systems in comparison to those estimated in the nucleus under well-controlled environment (Bentsen et al. 2012). Under these circumstances, the difference in the magnitude of the estimate was mainly due to scaling effects. Previous reviews (Nguyen 2016; Sae-Lim et al. 2016) also indicated that the genotype by environment (G×E) interaction was important when the production system differs markedly from the selection environment (e.g., fresh vs. brackish water), while the G×E was not significant when the two environments were similar. Even similar heritabilities in different environments do not necessary indicate no environment because they can result in different selection responses. A recent study in red tilapia reported significant impact of culture environments on genetic gain (Nguyen et al. 2017). Hence, the G×E effect should be examined on case by case basis to assist the design and conduct genetic improvement programs for aquaculture species.

### 5. Are the maternal and common environmental effects important?

The significant *c*^*2*^ effects for growth traits suggest that they should be included in statistical models to account for upward bias in genetic parameter estimates (Joshi et al. 2018). They also should be included in genetic evaluation systems to avoid any possible overestimation of breeding values as demonstrated in several studies (Hung et al. 2013a; Oliveira et al. 2016). To reduce the *c*^*2*^ effects in genetic improvement programs, early communal rearing of all families should be practised as soon as after birth. This approach can be implemented by using DNA markers (microsatellite or single nucleotide polymorphism, SNP) for genetic tagging and construction of pedigrees (Ninh et al. 2013; Whatmore et al. 2013).

### 6. Is the sample size sufficient to detect significant heritability?

Except for some studies where sample sizes were small (and the results were excluded from the present meta-analysis), the majority of reports included in this review had sufficient statistical power to detect significant heritability (h^2^) for traits with moderate to high heritability (i.e. growth traits) across aquaculture populations. The non-significant weighted h^2^ for other traits (flesh quality, fitness, reproduction) indicates that a much larger sample size and deeper pedigree are needed to obtain reliable genetic parameter estimates for low heritability traits; priority should be given to estimate genetic properties of these traits in future studies. Moreover, due to reproductive biology of many aquaculture species where synchronised mating is still difficult, *in-vitro* fertilisation in combination with advanced mating design such as factorial mating (Dupont-Nivet et al. 2006) should be applied to enable the estimation of different variance components; this would allow separation of additive genetics from non-additive genetic effects that are not widely understood in many fishes, crustaceans and mollucs. Across species, the conventional quantitative genetic (infinitesimal) model and family structure of aquaculture pedigrees did not allow a neat estimation of non-genetic components (maternal and common environmental effects), but they could be important for fitness-related traits and worth considering in future studies.

### 7. Were the heritabilities for important traits affected by publication bias?

The effect of publication bias on the heritability, for example, of body weight or morphological deformity was not significant in this study (see the Begger’s funnel plot and Egger regression asymmetry test in Supplementary File No.3). In addition, results of the sensitivity analysis performed by excluding studies with *z*-value greater than 3 (one study at a time or all together) gave almost identical heritability estimates for economically important traits (results not shown). This is also indicated by the narrow confidence interval, especially for body weight where the sample size for this trait was large. On the other hand, the effect of publication bias was present for other traits (flesh quality attributes and reproduction), partially due to the limited sample size reported for these characters.

### 8. Can selection for high performance also improve other traits?

With the current knowledge, there is a good understanding of the genetic associations among body traits (weight, length, width and depth) and those of growth traits with carcass yield, suggesting that either weight or length can be used as alternative selection criterion, and that selection for one of the two traits can simultaneously improve overall production performance of the animals (Hung and Nguyen 2014). Genetic correlations of body weight with carcass/flesh quality and reproductive traits are consistent with realised correlated changes in both fishes (Hamzah et al. 2014a; Hamzah et al. 2014b) and shrimps (Hung et al. 2013b). These changes are desired from genetic improvement perspectives. However, selection for high growth did not improve fillet yield in tilapia (Nguyen et al. 2010a) or caused insignificant changes in survival or fecundity per unit of female weight (Hamzah et al. 2015; Hamzah et al. 2014a). When growth is not of interest, fillet yield can still be improved through sib or indirection selection by using various yield predictors (Prchal et al. 2018; Vu et al. 2019). Furthermore, there is evidence in at least five populations of freshwater species (carps, tilapias and prawns) that selection for high growth didn’t cause adverse impact on survival (Vu et al. 2017). Based on the weighted genetic correlation estimates, it can be predicted that selection for high growth may not improve disease resistance across aquatic species, although the positive genetic correlation estimates between the two traits were reported in some studies. Furthermore, possible changes in fitness, social interaction or behavioural traits consequent to selection for high productivity were not well understood, because majority of breeding programs for aquaculture species have started rather recently, except for the two long run programs in Atlantic salmon in Norway (Gjedrem 2010) and Nile tilapia in the Philippines and then Malaysia (Hamzah et al. 2014a; Ponzoni et al. 2011).

### 9. Can traits showing antagonistic genetic relationships be improved simultaneously?

A number of traits included in this review showed antagonistic relationships that are not desired. For instance, based on the weighted genetic correlation estimate between body weight and sexual maturity, it can be predicted that selection for increased harvest body weight may produce early-maturing fish. On the one hand, the concomitant changes in the early maturity are desired for newly domesticated species in captivity (e.g., marine fishes). On the other, precocious reproduction in tilapia leads to low growth performance of stocked fish as a consequence of over-crowding and feed competition in production ponds. Under these circumstances, a restricted (or desired) selection index can be applied to simultaneously improve both traits (Nguyen et al. 2016; Sae-Lim et al. 2012). Multi-trait selection (Janssen et al. 2017) is recommended to maximise productivity and revenue as well as sustain long-term genetic improvement in order to meet future demand for high quality seafood by consumers and to adapt to environmental challenges.

### 10. Limitations and gaps in the literature

Only a few reports on heritability were found for traits that are expensive or difficult to measure, such as flesh/eating quality, behaviour and adaptive traits, across aquaculture species. The genetic association of these new traits with growth performance is inconclusive as the correlation estimates are not sufficient to support a systematic meta-analysis. To gain reliable estimates and better understanding about genetic relationships among these traits, a larger sample size from multigenerational pedigreed populations is needed. Until the present, objective measurements of traits that require slaughter of the animals are still not common in selective breeding programs for aquaculture species. For example, the application of near or mid infra-red spectroscopy may reduce costs of analysing fillet fatty acids in fish (Nguyen et al. 2010b), or quantitative real time PCR could be used to quantify viral copy numbers as an alternative approach to study disease resistance in shrimps (Phuthaworn et al. 2016). Challenges still remain to make objective and effective measurements of flesh/eating quality and behavioural/physiological traits in aquatic species. The availability of imaging technologies (X-ray, computed tomography CT) or machine vision systems also enables detailed examination of quality traits and deformities in fish at a reasonably low cost (Saberioon et al. 2016). In the near future, laboratory-generated data such as qPCR to quantify viral load/or titre level to select for disease resistance, and molecular genetic information or genome sequence, are expected to become available to study genetic architecture of new traits in aquatic species.

### 11. New prospects for understanding complex traits

In addition to the application of new measurements/data recording methods (section 4.8), the development of statistical approaches and algorithms such as restricted maximum likelihood method (Thompson 2008) applied to a multi-trait mixed model (Henderson 1975) helped to improve the accuracy of the genetic parameter estimates reported in the literature in comparison to the conventional analysis of variance (ANOVA) used 20-30 years ago. Recent expansions of the mixed model methodology have provided opportunities to study new traits, including social indirect genetic effects (Bijma 2014) or double hierarchical linear generalised model (Rönnegård and Lee 2013) to reduce environmental variation and improve uniformity (Hill and Mulder 2010) for aquatic species. The application of random regression analysis or cure survival model may help to understand disease tolerance and resistance in a range of species (Kause and Ødegård 2012). Structural equation models (Valente et al. 2010) can be applied to dissect complex biological relationships among traits, such as milk or fillet fatty acids (Bouwman et al. 2014). Particularly, the advent of high through-put whole genome sequencing and ‘omic’ related technologies coupled with better computational methods provide opportunities to better understand genetic architecture of quantitative complex traits in aquatic species. Genomic or genome-based selection has revolutionized breeding of dairy cattle and other livestock and plant sectors world-wide (Van Eenennaam et al. 2014). Recent studies in aquaculture species indicated that the accuracy of genomic prediction for quantitative traits was moderate to high (Nguyen et al. 2018b; Tsai et al. 2017; Yoshida et al. 2018), thereby the potential opportunities of genomic selection can offer to enhance aquaculture breeding in the near future.

## V. CONCLUSIONS

Meta-analysis of information concerning genetic parameters of complex quantitative traits in aquatic species revealed that:

1. The weighted means heritabilities (*h*^*2*^) for body and carcass traits in aquaculture species were moderate, whereas they were low and not significant for flesh quality attributes, except fillet fat content and fillet (or body) colour. Resistance against bacterial, viral and parasitic diseases was moderately heritable. The *h*^*2*^ estimates for fecundity and deformity were low and not significantly different from zero.
2. There were prospects to investigate new functional traits, namely behaviour, adaptation, uniformity, social interaction indirect genetic effects and immune response. There is very limited published information regarding genetic relationships between these novel characters and growth (or other traits) to enable multi-trait selection in future breeding programs.
3. The genetic correlations of body weight with colour and fillet fat content were moderate to high and positive. However, the estimates between growth and other flesh-quality attributes were weak and not significant. There was insignificant genetic association between growth and disease resistance. Genetic relationship of reproductive traits with body weight and growth-related characteristics merit further studies.
4. The maternal and common environmental effect (*c*^*2*^) accounted for 5 to 14% of the total phenotypic variance for growth-related traits, whereas it was not significant for flesh-quality attributes or reproductive characteristics, due to low number of investigations or non-optimal experimental design.
5. The effect of publication bias was not important for growth traits and deformity. On the other hand, it was apparent for disease resistance, flesh-quality attributes and reproductive characteristics.
6. There was small to moderate heterogeneity in the heritability and *c*^*2*^ effects for the majority of the traits studied. By contrast, the *Q*-statistics were significant for the genetic correlation estimates.
7. While results from the present meta-analysis provide fundamental information regarding genetic architecture of quantitative complex traits, genetic parameters should be estimated for each population before genetic improvement programs are initiated for a new species because the additive variation is different among populations within an individual species and it varies with environments and generations. The effect of genotype by environment is potentially important for complex traits, especially when the environments used in the nucleus differ from those practised under field production systems.

## VI. FUTURE DIRECTIONS

1. Future research should focus on novel characteristics that are emerging as potential selection criteria in genetic improvement programs, such as flesh/eating quality and new traits of commercial interest such as behaviour, adaptation, immune response and disease resistance.
2. Little is known about genetic relationships among these traits with body weight which justifies further studies.
3. Large sample size and multigenerational data from long term genetic improvement programs would increase the accuracy of genetic parameter estimates, especially for traits with low heritability (e.g., reproduction and deformity).
4. Laboratory based data, such as viral load quantified by qPCR and molecular genetic information or genome sequence should be increasingly incorporated into selective breeding programs for aquatic species
5. New methods of measurements should be applied to enable large-scale routine data collection for traits that are difficult or expensive to measure
6. Whole genome sequencing provides options to dissect the genetic architecture of quantitative complex traits across species.
7. Dissecting the quantitative genetic architecture of novel traits also requires development of new means of measurements to enable large-scale data recording as well as effective tools to manage pedigrees.

## VII. ACKNOWLEDGEMENTS

The author was funded by the Collaborative Research Network (CRN) from the Commonwealth Government of Australia and the University of the Sunshine Coast (USC).

## Supplementary files

Supplementary file No.1: PRISMA diagram flow to assess eligibility of studies included in the meta-analysis

Supplementary file No.2: A full list of references included in the review and analysis

Supplementary file No.3: Funnel plot to assess publication bias and sensitivity analyses

Supplementary file No.4: Detailed description of measurement methods for novel traits

## Glossary

**Quantitative complex traits** are influenced by both genetics and the environment, which can be expressed on continuous numerical scale (e.g. body weight) or non-continuous (meristic and threshold traits).

**Closed breeding nucleus** is a traditional breeding scheme where replacement animals come from the nucleus and no animals outside the nucleus are added to the population.

**Functional traits** are morphological, biochemical, physiological, structural, phenological or behavioural characteristics that influence organism performance or fitness.

**Heritability** estimates the degree of variation in a phenotypic trait in a population that is due to genetic variation between individuals in that population

**Common full-sib effects** are due to non-additive genetic, maternal and environmental effects and common family-tank/hapa effects, as a result of separate rearing of each family before physical tagging

**Phenotypic correlation** is a measure of the association between phenotypic values of characters

**Genetic correlation** is a measure of the association between breeding values of quantitative characters

**Random regression analysis** is a statistical method used to analyse repeated measurements over time or age (also termed as longitudinal data) to account for the mean and covariance structure that may change with time.

**Cure survival models** are used for modelling time-until-death data that include a fraction of non-affected individuals, i.e., those that are liable to die due to the infection.

**Structural equation models**, also known as path models, are used to understand the correlation structure of continuously variable data

**Double hierarchical linear generalised model (DHGLM)** is an extension of generalized linear model to include a structure for one or more variance components and/or the residual variance. The method is used for estimation in models with genetically structured heterogeneity of residual variance for large quantitative data sets

