## Supplementary Files for "A systematic review and meta-analysis of genetic parameters for complex quantitative traits in aquatic animal species"

### Supplementary file N.1: PRISMA 2009 Flow Diagram

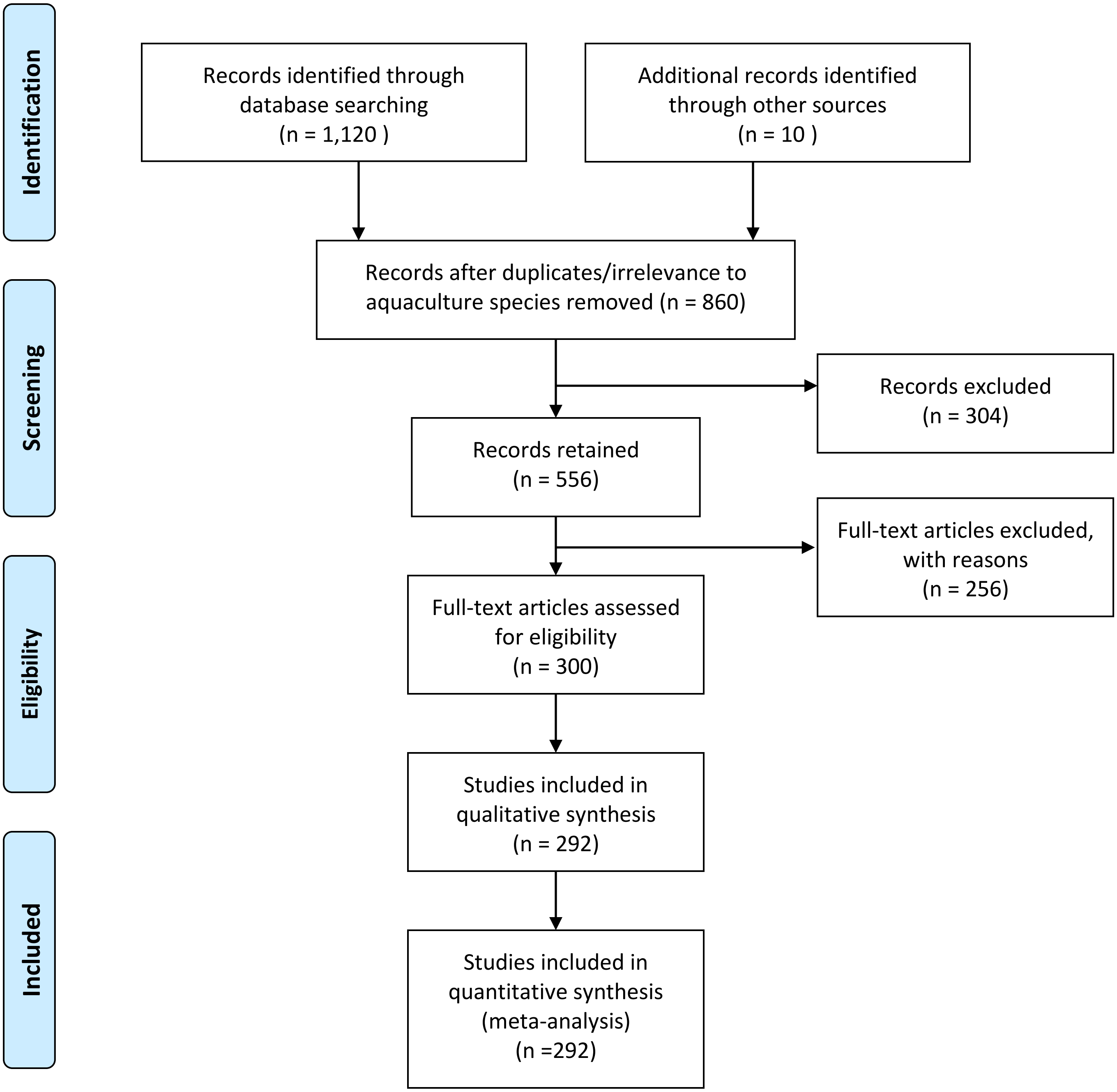

*From:* Moher D, Liberati A, Tetzlaff J, Altman DG, The PRISMA Group (2009). *P*referred *R*eporting *I*tems for *S*ystematic Reviews and *M*eta-*A*nalyses: The PRISMA Statement. PLoS Med 6(6): e1000097. doi:10.1371/journal.pmed1000097

**For more information, visit www.prisma-statement.org.**

**Supplementary file N.2: FULL LIST OF ARTICLES INCLUDED IN THE REVIEW**

Alcapán, A.C., Nespolo, R.F., Toro, J.E. (2007) Heritability of body size in the Chilean blue mussel (Mytilus chilensis Hupe 1854): effects of environment and ageing. *Aquaculture Research* **38,** 313-320.

Antonello, J., Massault, C., Franch, R.*, et al.* (2009) Estimates of heritability and genetic correlation for body length and resistance to fish pasteurellosis in the gilthead sea bream (Sparus aurata L.). *Aquaculture* **298,** 29-35.

Arcos, F.G., Racotta, I.S., Ibarra, A.M. (2004) Genetic parameter estimates for reproductive traits and egg composition in Pacific white shrimp *Penaeus (Litopenaeus) vannamei*. *Aquaculture* **236,** 151-165.

Argue, B.J., Arce, S.M., Lotz, J.M., Moss, S.M. (2002) Selective breeding of Pacific white shrimp *(Litopenaeus vannamei)* for growth and resistance to Taura Syndrome Virus. *Aquaculture* **204,** 447-460.

Azéma, P., Lamy, J.-B., Boudry, P., Renault, T., Travers, M.-A., Dégremont, L. (2017) Genetic parameters of resistance to Vibrio aestuarianus, and OsHV-1 infections in the Pacific oyster, Crassostrea gigas, at three different life stages. *Genetics Selection Evolution* **49,** 23.

Bangera, R., Correa, K., Lhorente, J.P., Figueroa, R., Yáñez, J.M. (2017) Genomic predictions can accelerate selection for resistance against Piscirickettsia salmonis in Atlantic salmon (Salmo salar). *BMC Genomics* **18,** 121.

Bangera, R., Ødegård, J., Mikkelsen, H.*, et al.* (2014) Genetic analysis of francisellosis field outbreak in Atlantic cod (Gadus morhua L.) using an ordinal threshold model. *Aquaculture* **420,** S50-S56.

Bangera, R., Ødegård, J., Nielsen, H., Gjøen, H., Mortensen, A. (2013) Genetic analysis of vibriosis and viral nervous necrosis resistance in Atlantic cod (L.) using a cure model. *Journal of Animal Science* **91,** 3574-3582.

Bangera, R., Ødegård, J., Præbel, A.K., Mortensen, A., Nielsen, H.M. (2011) Genetic correlations between growth rate and resistance to vibriosis and viral nervous necrosis in Atlantic cod (Gadus morhua L.). *Aquaculture* **317,** 67-73.

Bardon, A., Vandeputte, M., Dupont-Nivet, M.*, et al.* (2009) What is the heritable component of spinal deformities in the European sea bass (Dicentrarchus labrax)? *Aquaculture* **294,** 194-201.

Bentsen, H.B., Gjerde, B., Nguyen, N.H.*, et al.* (2012) Genetic improvement of farmed tilapias: Genetic parameters for body weight at harvest in Nile tilapia (Oreochromis niloticus) during five generations of testing in multiple environments. *Aquaculture* **338,** 56-65.

Benzie, J.A.H., Kenway, M., Trott, L. (1997) Estimates for the heritability of size in juvenile *Penaeus monodon* prawns from half-sib matings. *Aquaculture* **152,** 49-53.

Blonk, R., Komen, J., Tenghe, A., Kamstra, A., Van Arendonk, J. (2010a) Heritability of shape in common sole, Solea solea, estimated from image analysis data. *Aquaculture* **307,** 6-11.

Blonk, R.J., Komen, H., Kamstra, A., van Arendonk, J.A. (2010b) Effects of grading on heritability estimates under commercial conditions: A case study with common sole, Solea solea. *Aquaculture* **300,** 43-49.

Bolivar, R.B., Newkirk, G.F. (2002) Response to within family selection for body weight in Nile tilapia (*Oreochromis niloticus*) using a single-trait animal model. *Aquaculture* **204,** 371-381.

Bosworth, B., Waldbieser, G. (2014) General and specific combining ability of male blue catfish (Ictalurus furcatus) and female channel catfish (Ictalurus punctatus) for growth and carcass yield of their F 1 hybrid progeny. *Aquaculture* **420,** 147-153.

Boudry, P., Collet, B., Cornette, F., Hervouet, V., Bonhomme, F. (2002) High variance in reproductive success of the Pacific oyster (*Crassostrea gigas*, Thunberg) revealed by microsatellite-based parentage analysis of multifactorial crosses. *Aquaculture* **204,** 283-296.

Boudry, P., Dégremont, L., Taris, N., McCombie, H., Haffray, P., Ernande, B. (2004) Genetic variability and selective breeding for traits of aquacultural interest in the Pacific oyster (Crassostrea gigas). *Bulletin of the Aquaculture Association of Canada* **104,** 12-18.

Brokordt, K., Farías, W., Lhorente, J.P., Winkler, F. (2012) Heritability and genetic correlations of escape behaviours in juvenile scallop Argopecten purpuratus. *Animal Behaviour* **84,** 479-484.

Brokordt, K.B., Winkler, F.M., Farías, W.J.*, et al.* (2015) Changes of heritability and genetic correlations in production traits over time in red abalone (Haliotis rufescens) under culture. *Aquaculture Research* **46,** 2248-2259.

Caballero-Zamora, A., Montaldo, H., Campos-Montes, G., Cienfuegos-Rivas, E., Martínez-Ortega, A., Castillo-Juárez, H. Estimation of (co) variance components for body weight and survival in the presence of a White Spot Syndrome Virus (WSSV) natural outbreak in the Pacific White Shrimp Penaeus (Litopenaeus) vannamei. (Proceedings of the 10th World Congress on Genetics Applied to Livestock Production, 2014). Asas, City.

Caballero-Zamora, A., Cienfuegos-Rivas, E.G., Montaldo, H.H., Campos-Montes, G.R., Martínez-Ortega, A., Castillo-Juárez, H. (2015) Genetic parameters for spawning and growth traits in the Pacific white shrimp (Penaeus (Litopenaeus) vannamei). *Aquaculture Research* **46,** 833-839.

Camara, M.D., Yen, S., Kaspar, H.F.*, et al.* (2017) Assessment of heat shock and laboratory virus challenges to selectively breed for ostreid herpesvirus 1 (OsHV-1) resistance in the Pacific oyster, Crassostrea gigas. *Aquaculture* **469,** 50-58.

Campos-Montes, G.R., Montaldo, H.H., Armenta-Córdova, M., Martínez-Ortega, A., Caballero-Zamora, A., Castillo-Juárez, H. (2017) Incorporation of tail weight and tail percentage at harvest size in selection programs for the Pacific white shrimp Penaeus (Litopenaeus) vannamei. *Aquaculture* **468,** 293-296.

Campos-Montes, G.R., Montaldo, H.H., Martínez-Ortega, A., Jiménez, A.M., Castillo-Juárez, H. (2013) Genetic parameters for growth and survival traits in Pacific white shrimp Penaeus (Litopenaeus) vannamei from a nucleus population undergoing a two-stage selection program. *Aquaculture International* **21,** 299-310.

Cao, X., Wang, H., Yao, H.*, et al.* (2012) Evaluation of 1-stage and 2-stage selection in yellow perch I: Genetic and phenotypic parameters for body weight of F fish reared in ponds using microsatellite parentage assignment. *Journal of Animal Science* **90,** 27-36.

Castillo-Juárez, H., Casares, J.C.Q., Campos-Montes, G., Villela, C.C., Ortega, A.M., Montaldo, H.H. (2007) Heritability for body weight at harvest size in the Pacific white shrimp, *Penaeus (Litopenaeus) vannamei,* from a multi-environment experiment using univariate and multivariate animal models. *Aquaculture* **273,** 42-49.

Charo-Karisa, H., Bovenhuis, H., Rezk, M.A., Ponzoni, R.W., van Arendonk, J.A.M., Komen, H. (2007) Phenotypic and genetic parameters for body measurements, reproductive traits and gut length of Nile tilapia (*Oreochromis niloticus*) selected for growth in low-input earthen ponds. *Aquaculture* **273,** 15-23.

Charo-Karisa, H., Komen, H., Rezk, M., Ponzoni, R., van Arendonk, J., Bovenhuis, H. (2006) Heritability estimates and response to selection for growth of Nile tilapia (Oreochromis niloticus) in low-input earthen ponds. *Aquaculture* **261,** 479 - 486.

Charo-Karisa, H., Rezk, M.A., Bovenhuis, H., Komen, H. (2005) Heritability of cold tolerance in Nile tilapia, Oreochromis niloticus, juveniles. *Aquaculture* **249,** 115-123.

Cheryl, D., Antti, K., Juha, K. (2007) Breeding salmonids for feed efficiency in current fishmeal and future plant-based diet environments. *Genet. Sel. Evol* **39,** 431-446.

Chevassus, B., Quillet, E., Krieg, F.*, et al.* (2004) Enhanced individual selection for selecting fast growing fish: the “PROSPER” method, with application on brown trout (Salmo trutta fario). *Genetics Selection Evolution* **36,** 643-661.

Chiasson, M., Quinton, C., Danzmann, R., Ferguson, M. (2013) Comparative analysis of genetic parameters and quantitative trait loci for growth traits in Fraser strain Arctic charr () reared in freshwater and brackish water environments. *Journal of Animal Science* **91,** 2047-2056.

de Melo, C.M.R., Durland, E., Langdon, C. (2016) Improvements in desirable traits of the Pacific oyster, Crassostrea gigas, as a result of five generations of selection on the West Coast, USA. *Aquaculture* **460,** 105-115.

De Verdal, H., Rosario, W., Vandeputte, M.*, et al.* (2014) Response to selection for growth in an interspecific hybrid between Oreochromis mossambicus and O. niloticus in two distinct environments. *Aquaculture* **430,** 159-165.

Dégremont, L., Bédier, E., Boudry, P. (2010) Summer mortality of hatchery-produced Pacific oyster spat (Crassostrea gigas). II. Response to selection for survival and its influence on growth and yield. *Aquaculture* **299,** 21-29.

Dégremont, L., Ernande, B., Bédier, E., Boudry, P. (2007) Summer mortality of hatchery-produced Pacific oyster spat (Crassostrea gigas). I. Estimation of genetic parameters for survival and growth. *Aquaculture* **262,** 41-53.

Dégremont, L., Garcia, C., Allen, S.K. (2015) Genetic improvement for disease resistance in oysters: a review. *Journal of Invertebrate Pathology* **131,** 226-241.

Deng, Y., Fu, S., Du, X., Wang, Q. (2009) Realized heritability and genetic gain estimates of larval shell length in the Chinese pearl oyster Pinctada martensii at three different salinities. *North American Journal of Aquaculture* **71,** 302-306.

Difford, G., Vlok, A., Rhode, C., Brink, D. (2017) Heritability of growth traits in South African Abalone (Haliotis midae L.) using the 'internal reference'method. *Aquaculture* **468,** 451-457.

Domingos, J.A., Smith-Keune, C., Robinson, N., Loughnan, S., Harrison, P., Jerry, D.R. (2013) Heritability of harvest growth traits and genotype–environment interactions in barramundi, *Lates calcarifer* (Bloch). *Aquaculture* **402–403,** 66-75.

Dong, Z., Nguyen, N.H., Zhu, W. (2015) Genetic evaluation of a selective breeding program for common carp *Cyprinus carpio* conducted from 2004 to 2014. *BMC Genetics* **16,** 1-9.

Drangsholt, T., Gjerde, B., Ødegård, J., Finne-Fridell, F., Evensen, Ø., Bentsen, H. (2011) Quantitative genetics of disease resistance in vaccinated and unvaccinated Atlantic salmon (Salmo salar L.). *Heredity* **107,** 471-477.

Drangsholt, T.M.K., Damsgård, B., Olesen, I. (2014) Quantitative genetics of behavioral responsiveness in Atlantic cod (Gadus morhua L.). *Aquaculture* **420–421,** 282-287.

Drangsholt, T.M.K., Gjerde, B., Ødegård, J., Finne-Fridell, F., Evensen, Ø., Bentsen, H.B. (2012) Genetic correlations between disease resistance, vaccine-induced side effects and harvest body weight in Atlantic salmon (Salmo salar). *Aquaculture* **324,** 312-314.

Dufflocq, P., Lhorente, J.P., Bangera, R., Neira, R., Newman, S., Yáñez, J.M. (2016) Correlated response of flesh color to selection for harvest weight in coho salmon (Oncorhynchus kisutch). *Aquaculture*.

Dupont-Nivet, M., Karahan-Nomm, B., Vergnet, A.*, et al.* (2010) Genotype by environment interactions for growth in European seabass (Dicentrarchus labrax) are large when growth rate rather than weight is considered. *Aquaculture* **306,** 365-368.

Dupont-Nivet, M., Vandeputte, M., Vergnet, A.*, et al.* (2008) Heritabilities and GxE interactions for growth in the European sea bass (*Dicentrarchus labrax* L.) using a marker-based pedigree. *Aquaculture* **275,** 81-87.

Eknath, A.E., Bentsen, H.B., Ponzoni, R.W.*, et al.* (2007) Genetic improvement of farmed tilapias: Composition and genetic parameters of a synthetic base population of *Oreochromis niloticus* for selective breeding. *Aquaculture* **273,** 1-14.

Evans, S., Camara, M.D., Langdon, C.J. (2009) Heritability of shell pigmentation in the Pacific oyster, Crassostrea gigas. *Aquaculture* **286,** 211-216.

Evans, S., Langdon, C. (2006) Effects of genotype × environment interactions on the selection of broadly adapted Pacific oysters (*Crassostrea gigas*). *Aquaculture* **261,** 522-534.

Evenhuis, J., Leeds, T., Marancik, D., LaPatra, S., Wiens, G. (2015) Rainbow trout () resistance to columnaris disease is heritable and favorably correlated with bacterial cold water disease resistance. *Journal of Animal Science* **93,** 1546-1554.

Ferrari, S., Horri, K., Allal, F.*, et al.* (2016) Heritability of Boldness and Hypoxia Avoidance in European Seabass, Dicentrarchus labrax. *PloS One* **11,** e0168506.

Fishback, A.G., Danzmann, R.G., Ferguson, M.M., Gibson, J.P. (2002) Estimates of genetic parameters and genotype by environment interactions for growth traits of rainbow trout (*Oncorhynchus mykiss*) as inferred using molecular pedigrees. *Aquaculture* **206,** 137-150.

Fjalestad, K.T., Gjedrem, T., Gjerde, B. (1993) Genetic improvement of disease resistance in fish: an overview. *Aquaculture* **111,** 65-74.

Fjelldal, P., Hansen, T., Breck, O.*, et al.* (2009) Supplementation of dietary minerals during the early seawater phase increase vertebral strength and reduce the prevalence of vertebral deformities in fast-growing under-yearling Atlantic salmon (Salmo salar L.) smolt. *Aquaculture Nutrition* **15,** 366-378.

Gall, G.A., Bakar, Y. (1999) Stocking density and tank size in the design of breed improvement programs for body size of tilapia. *Aquaculture* **173,** 197-205.

Gall, G.A., Bakar, Y. (2002) Application of mixed-model techniques to fish breed improvement: analysis of breeding-value selection to increase 98-day body weight in tilapia. *Aquaculture* **212,** 93-113.

Gall, G.A., Neira, R. (2004) Genetic analysis of female reproduction traits of farmed coho salmon (Oncorhyncus kisutch). *Aquaculture* **234,** 143-154.

Gallardo, J.A., Lhorente, J.P., Neira, R. (2010) The consequences of including non-additive effects on the genetic evaluation of harvest body weight in Coho salmon (Oncorhynchus kisutch). *Genetics Selection Evolution* **42,** 19.

Garber, A.F., Tosh, J.J., Fordham, S.E.*, et al.* (2010) Survival and growth traits at harvest of communally reared families of Atlantic cod (Gadus morhua). *Aquaculture* **307,** 12-19.

García-Celdrán, M., Cutáková, Z., Ramis, G.*, et al.* (2016) Estimates of heritabilities and genetic correlations of skeletal deformities and uninflated swimbladder in a reared gilthead sea bream (Sparus aurata L.) juvenile population sourced from three broodstocks along the Spanish coasts. *Aquaculture* **464,** 601-608.

García-Celdrán, M., Ramis, G., Manchado, M., Estévez, A., Navarro, A., Armero, E. (2015) Estimates of heritabilities and genetic correlations of raw flesh quality traits in a reared gilthead sea bream (Sparus aurata L.) population sourced from broodstocks along the Spanish coasts. *Aquaculture* **446,** 181-186.

Gheyas, A.A., Woolliams, J.A., Taggart, J.B.*, et al.* (2009) Heritability estimation of silver carp (*Hypophthalmichthys molitrix*) harvest traits using microsatellite based parentage assignment. *Aquaculture* **294,** 187-193.

Gima, M.E., Gima, A., Hutson, A.*, et al.* (2014) Realized heritability and response to selection for fecundity, hatching rate and fry/Kg for channel catfish females (Ictalurus punctatus) induced to ovulate and fertilized with blue catfish (Ictalurus furcatus) males for the production of hybrid catfish embryos. *Aquaculture* **420,** S36-S41.

Gitterle, T., Gjerde, B., Cock, J.*, et al.* (2006) Optimization of experimental infection protocols for the estimation of genetic parameters of resistance to White Spot Syndrome Virus (WSSV) in *Penaeus (Litopenaeus) vannamei*. *Aquaculture* **261,** 501-509.

Gitterle, T., Rye, M., Salte, R.*, et al.* (2005a) Genetic (co)variation in harvest body weight and survival in *Penaeus (Litopenaeus) vannamei* under standard commercial conditions. *Aquaculture* **243,** 83-92.

Gitterle, T., Salte, R., Gjerde, B.*, et al.* (2005b) Genetic (co)variation in resistance to White Spot Syndrome Virus (WSSV) and harvest weight in *Penaeus (Litopenaeus) vannamei*. *Aquaculture* **246,** 139-149.

Gjedrem, T. (1997) Flesh quality improvement in fish through breeding. *Aquaculture International* **5,** 197-206.
Gjedrem, T. (2000) Genetic improvement of cold-water fish species. *Aquaculture Research* **31,** 25-33.
Gjerde, B., Evensen, Ø., Bentsen, H.B., Storset, A. (2009) Genetic (co) variation of vaccine injuries and innate resistance to furunculosis (Aeromonas salmonicida) and infectious salmon anaemia (ISA) in Atlantic salmon (Salmo salar). *Aquaculture* **287,** 52-58.
Gjerde, B., Mengistu, S.B., Ødegård, J., Johansen, H., Altamirano, D.S. (2012) Quantitative genetics of body weight, fillet weight and fillet yield in Nile tilapia (Oreochromis niloticus). *Aquaculture* **342–343,** 117-124.
Gjerde, B., Ødegård, J., Thorland, I. (2011) Estimates of genetic variation in the susceptibility of Atlantic salmon (Salmo salar) to the salmon louse Lepeophtheirus salmonis. *Aquaculture* **314,** 66-72.
Gjerde, B., Pante, M.J.R., Baeverfjord, G. (2005) Genetic variation for a vertebral deformity in Atlantic salmon (Salmo salar). *Aquaculture* **244,** 77-87.
Gjerde, B., Schaeffer, L. (1989) Body traits in rainbow trout. 2. Estimates of heritabilities and of phenotypic and genetic correlations. *Aquaculture* **80,** 25 - 44.
Gjerde, B., Simianer, H., Refstie, T. (1994) Estimates of genetic and phenotypic parameters for body weight, growth rate and sexual maturity in Atlantic salmon. *Livestock Production Science* **38,** 133-143.
Gjerde, B., Terjesen, B.F., Barr, Y., Lein, I., Thorland, I. (2004) Genetic variation for juvenile growth and survival in Atlantic cod (*Gadus morhua*). *Aquaculture* **236,** 167-177.
Gjøen, H., Bentsen, H. (1997) Past, present, and future of genetic improvement in salmon aquaculture. *ICES Journal of Marine Science: Journal du Conseil* **54,** 1009-1014.
Gjøen, H.M., Refstie, T., Ulla, O., Gjerde, B. (1997) Genetic correlations between survival of Atlantic salmon in challenge and field tests. *Aquaculture* **158,** 277-288.
Glover, K., Aasmundstad, T., Nilsen, F., Storset, A., Skaala, Ø. (2005) Variation of Atlantic salmon families (Salmo salar L.) in susceptibility to the sea lice Lepeophtheirus salmonis and Caligus elongatus. *Aquaculture* **245,** 19-30.
Gopal, C., Gopikrishna, G., Krishna, G*., et al.* (2010) Weight and time of onset of female-superior sexual dimorphism in pond reared Penaeus monodon. *Aquaculture* **300,** 237-239.
Grima, L., Chatain, B., Ruelle, F*., et al.* (2010) In search for indirect criteria to improve feed utilization efficiency in sea bass (Dicentrarchus labrax): Part II: Heritability of weight loss during feed deprivation and weight gain during re-feeding periods. *Aquaculture* **302,** 169 - 174.
Guan, J., Hu, Y., Wang, M., Wang, W., Kong, J., Luan, S. (2016) Estimating genetic parameters and genotype-by-environment interactions in body traits of turbot in two different rearing environments. *Aquaculture* **450,** 321-327.
Guiñez, R., Toro, J.E., Krapivka, S., Alcapán, A.C., Oyarzún, P.A. (2016) Heritabilities and genetic correlation of shell thickness and shell length growth in a mussel, Mytilus chilensis (Bivalvia: Mytilidae). *Aquaculture Research*.
Guy, D.R., Bishop, S.C., Brotherstone, S*., et al.* (2006) Analysis of the incidence of infectious pancreatic necrosis mortality in pedigreed Atlantic salmon, Salmo salar L., populations. *Journal of Fish Diseases* **29,** 637-647.
Guy, D.R., Bishop, S.C., Woolliams, J.A., Brotherstone, S. (2009) Genetic parameters for resistance to Infectious Pancreatic Necrosis in pedigreed Atlantic salmon (*Salmo salar*) post-smolts using a Reduced Animal Model. *Aquaculture* **290,** 229-235.
Haffray, P., Bugeon, J., Pincent, C*., et al.* (2012) Negative genetic correlations between production traits and head or bony tissues in large all-female rainbow trout (Oncorhynchus mykiss). *Aquaculture* **368,** 145-152.
Haffray, P., Bugeon, J., Rivard, Q*., et al.* (2013) Genetic parameters of in-vivo prediction of carcass, head and fillet yields by internal ultrasound and 2D external imagery in large rainbow trout (Oncorhynchus mykiss). *Aquaculture* **410,** 236-244.

Hamzah, A., Mekkawy, W., Khaw, H.L., *et al.* (2015) Genetic parameters for survival during the grow-out period in the GIFT strain of Nile tilapia (Oreochromis niloticus) and correlated response to selection for harvest weight. *Aquaculture Research*, doi: 10.1111/are.12859.
Hamzah, A., Nguyen, N.H., Mekkawy, W., *et al.* (2014a) Genetic parameters and correlated responses in female reproductive traits in the GIFT strain. *Aquaculture Research*, doi: 10.1111/are.12608.
Hamzah, A., Nguyen, N.H., Mekkawy, W., *et al.* (2014b) Flesh characteristics: estimation of genetic parameters and correlated responses to selection for growth rate in the GIFT strain. *Aquaculture Research*, doi: 10.1111/are.12667.
Hamzah, A., Ponzoni, R.W., Nguyen, N.H., Khaw, H.L., Yee, H.Y., Nor, S.A.M. (2014c) Genetic evaluation of the Genetically Improved Farmed Tilapia (GIFT) strain over ten generations of selection in Malaysia. *Journal of Tropical Agricultural Science* **37,** 411-429.
He, J., Gao, H., Xu, P., Yang, R. (2015) Genetic parameters for different growth scales in GIFT strain of Nile tilapia (Oreochromis niloticus). *Journal of Animal Breeding and Genetics*, n/a-n/a.
He, M., Guan, Y., Yuan, T., Zhang, H. (2008) Realized heritability and response to selection for shell height in the pearl oyster Pinctada fucata (Gould). *Aquaculture Research* **39,** 801-805.
Henryon, M., Berg, P., Olesen, N.J., *et al.* (2005) Selective breeding provides an approach to increase resistance of rainbow trout (*Onchorhynchus mykiss*) to the diseases, enteric redmouth disease, rainbow trout fry syndrome, and viral haemorrhagic septicaemia. *Aquaculture* **250,** 621-636.
Henryon, M., Jokumsen, A., Berg, P., *et al.* (2002) Genetic variation for growth rate, feed conversion efficiency, and disease resistance exists within a farmed population of rainbow trout. *Aquaculture* **209,** 59 - 76.
Hung, D., Nguyen, H.N. (2014) Genetic inheritance of female and male morphotypes in giant freshwater prawn *Macrobrachium rosenbergii*. *PloS One* **9,** e90142.
Hung, D., Nguyen, N.H., Hurwood, D.A., Mather, P.B. (2014) Quantitative genetic parameters for body traits at different ages in a cultured stock of giant freshwater prawn (*Macrobrachium rosenbergii*) selected for fast growth. *Marine and Freshwater Research* **65,** 198-205.
Hung, D., Nguyen, N.H., Ponzoni, R.W., Hurwood, D.A., Mather, P.B. (2013) Quantitative genetic parameter estimates for body and carcass traits in a cultured stock of giant freshwater prawn (Macrobrachium rosenbergii) selected for harvest weight in Vietnam. *Aquaculture* **404–405,** 122-129.
Hung, D., Nguyen., N.H. (2014) Genetic inheritance of female and male morphotypes in Giant freshwater prawn *Macrobrachium roseinbergii*. *PloS One* **9,** e90142.
Ibarra, A., Famula, T. (2008) Genotype by environment interaction for adult body weights of shrimp *Penaeus vannamei* when grown at low and high densitie. *Genetics Selection Evolution* **40,** 541 - 551.
Ibarra, A.M., Ramirez, J.L., Ruiz, C.A., Cruz, P., Avila, S. (1999) Realized heritabilities and genetic correlation after dual selection for total weight and shell width in catarina scallop (Argopecten ventricosus). *Aquaculture* **175,** 227-241.
Iwamoto, R., Myers, J., Hershberger, W. (1986) Genotype-environment interactions for growth of rainbow trout, Salmo gairdneri. *Aquaculture* **57,** 153-161.
Iwamoto, R., Myers, J., Hershberger, W. (1990) Heritability and genetic correlations for flesh coloration in pen-reared coho salmon. *Aquaculture* **86,** 181-190.
Janhunen, M., Kause, A., Vehviläinen, H., Järvisalo, O. (2012) Genetics of microenvironmental sensitivity of body weight in rainbow trout (Oncorhynchus mykiss) selected for improved growth. *PloS One* **7,** e38766.
Janhunen, M., Kause, A., Vehviläinen, H., Nousiainen, A., Koskinen, H. (2013) Accounting for early rearing density effects on growth in the genetic evaluation of rainbow trout. *Journal of Animal Science* **91,** 5144-5152.

Janhunen, M., Koskela, J., Ninh, N.H.*, et al.* (2016) Thermal sensitivity of growth indicates heritable variation in 1-year-old rainbow trout (Oncorhynchus mykiss). *Genetics Selection Evolution* **48,** 94.

Jarayabhand, P., Thavornyutikarn, M. (1995) Realized heritability estimation on growth rate of oyster, Saccostrea cucullata Born, 1778. *Aquaculture* **138,** 111-118.

Jerry, D.R., Kvingedal, R., Lind, C.E., Evans, B.S., Taylor, J.J., Safari, A.E. (2012) Donor-oyster derived heritability estimates and the effect of genotype× environment interaction on the production of pearl quality traits in the silver-lip pearl oyster, Pinctada maxima. *Aquaculture* **338,** 66-71.

Kahar, S., Debes, P.V., Vuori, K.A., Vähä, J.-P., Vasemägi, A. (2016) Heritability, environmental effects, and genetic and phenotypic correlations of oxidative stress resistance-related enzyme activities during early life stages in Atlantic salmon. *Evolutionary Biology* **43,** 215-226.

Karahan, B., Chatain, B., Chavanne, H.*, et al.* (2013) Heritabilities and correlations of deformities and growth-related traits in the European sea bass (Dicentrarchus labrax, L) in four different sites. *Aquaculture Research* **44,** 289-299.

Kause, A., Paananen, T., Ritola, O., Koskinen, H. (2007) Direct and indirect selection of visceral lipid weight, fillet weight, and fillet percentage in a rainbow trout breeding program. *Journal of Animal Science* **85,** 3218-3227.

Kause, A., Quinton, C., Airaksinen, S., Ruohonen, K., Koskela, J. (2011) Quality and production trait genetics of farmed European whitefish. *Journal of Animal Science* **89,** 959-971.

Kause, A., Ritola, O., Paananen, T., Mäntysaari, E., Eskelinen, U. (2002) Coupling body weight and its composition: a quantitative genetic analysis in rainbow trout. *Aquaculture* **211,** 65-79.

Kause, A., Ritola, O., Paananen, T., Mäntysaari, E., Eskelinen, U. (2003) Selection against early maturity in large rainbow trout *Oncorhynchus mykiss*: the quantitative genetics of sexual dimorphism and genotype-by-environment interactions. *Aquaculture* **228,** 53-68.

Kause, A., Ritola, O., Paananen, T., Wahlroos, H., Mäntysaari, E.A. (2005) Genetic trends in growth, sexual maturity and skeletal deformations, and rate of inbreeding in a breeding programme for rainbow trout (*Oncorhynchus mykiss*). *Aquaculture* **247,** 177-187.

Kause, A., Stien, L., Rungruangsak-Torrissen, K., Ritola, O., Ruohonen, K., Kiessling, A. (2008) Image analysis as a tool to facilitate selective breeding of quality traits in rainbow trout. *Livestock Science* **114,** 315-324.

Kause, A., Tobin, D., Houlihan, D.F.*, et al.* (2006) Feed efficiency of rainbow trout can be improved through selection: Different genetic potential on alternative diets. *Journal of Animal Science* **84,** 807-817.

Kenway, M., Macbeth, M., Salmon, M.*, et al.* (2006) Heritability and genetic correlations of growth and survival in black tiger prawn *Penaeus monodon* reared in tanks. *Aquaculture* **259,** 138-145.

Kettunen, A., Fjalestad, K.T. (2007) Genetic parameters for important traits in the breeding program for Atlantic cod (Gadus morhua L.). *Aquaculture* **272,** S276-S276.

Khaw, H.L., Bovenhuis, H., Ponzoni, R.W., Rezk, M.A., Charo-Karisa, H., Komen, H. (2009) Genetic analysis of Nile tilapia (*Oreochromis niloticus*) selection line reared in two input environments. *Aquaculture* **294,** 37-42.

Khaw, H.L., Ponzoni, R.W., Hamzah, A., Abu-Bakar, K.R., Bijma, P. (2012) Genotype by production environment interaction in the GIFT strain of Nile tilapia (*Oreochromis niloticus*). *Aquaculture* **326–329,** 53-60.

Khaw, H.L., Ponzoni, R.W., Yee, H.Y., bin Aziz, M.A., Bijma, P. (2016a) Genetic and non-genetic indirect effects for harvest weight in the GIFT strain of Nile tilapia (Oreochromis niloticus). *Aquaculture* **450,** 154-161.

Khaw, H.L., Ponzoni, R.W., Yee, H.Y.*, et al.* (2016b) Genetic variance for uniformity of harvest weight in Nile tilapia (Oreochromis niloticus). *Aquaculture* **451,** 113-120.

Kitcharoen, N., Rungsin, W., Koonawootrittriron, S., Na-Nakorn, U. (2012) Heritability for growth traits in giant freshwater prawn, Macrobrachium rosenbergii (de Mann 1879) based on best linear unbiased prediction methodology. *Aquaculture Research* **43,** 19-25.

Kjøglum, S., Henryon, M., Aasmundstad, T., Korsgaard, I. (2008) Selective breeding can increase resistance of Atlantic salmon to furunculosis, infectious salmon anaemia and infectious pancreatic necrosis. *Aquaculture Research* **39,** 498-505.

Knibb, W., Miller, A., Quinn, J., D'Antignana, T., Nguyen, N.H. (2016) Comparison of lines shows selection response in kingfish (Seriola lalandi). *Aquaculture* **452,** 318-325.

Kocour, M., Linhart, O., Gela, D., Rodina, M. (2005) Growth Performance of All-Female and Mixed-Sex Common Carp Cyprinus Carpio L. Populations in the Central Europe Climatic Conditions. *Journal of the World Aquaculture Society* **36,** 103-113.

Kocour, M., Linhart, O., Vandeputte, M. (2006) Mouth and fin deformities in common carp: is there a genetic basis? *Aquaculture Research* **37,** 419-422.

Kocour, M., Mauger, S., Rodina, M., Gela, D., Linhart, O., Vandeputte, M. (2007) Heritability estimates for processing and quality traits in common carp (Cyprinus carpio L.) using a molecular pedigree. *Aquaculture* **270,** 43-50.

Kolstad, K., Heuch, P.A., Gjerde, B., Gjedrem, T., Salte, R. (2005) Genetic variation in resistance of Atlantic salmon (Salmo salar) to the salmon louse Lepeophtheirus salmonis. *Aquaculture* **247**.

Kolstad, K., Thorland, I., Refstie, T., Gjerde, B. (2006) Genetic variation and genotype by location interaction in body weight, spinal deformity and sexual maturity in Atlantic cod (*Gadus morhua*) reared at different locations off Norway. *Aquaculture* **259,** 66-73.

Kong, N., Li, Q., Yu, H., Kong, L.F. (2015) Heritability estimates for growth-related traits in the Pacific oyster (Crassostrea gigas) using a molecular pedigree. *Aquaculture Research* **46,** 499-508.

Kortet, R., Vainikka, A., Janhunen, M., Piironen, J., Hyvärinen, P. (2014) Behavioral variation shows heritability in juvenile brown trout Salmo trutta. *Behavioral ecology and sociobiology* **68,** 927-934.

Krishna, G., Gopikrishna, G., Gopal, C., *et al.* (2011) Genetic parameters for growth and survival in Penaeus monodon cultured in India. *Aquaculture* **318,** 74-78.

Kristjánsson, T., Arnason, T. (2014) Heritability of economically important traits in the Atlantic cod Gadus morhua L. *Aquaculture Research*.

Kronert, U., Hörstgen-Schwark, G., Langholz, H.-J. (1989) Prospects of selecting for late maturity in tilapia (Oreochromis niloticus): I. Family studies under laboratory conditions. *Aquaculture* **77,** 113-121.

Kube, P.D., Taylor, R.S., Elliott, N.G. (2012) Genetic variation in parasite resistance of Atlantic salmon to amoebic gill disease over multiple infections. *Aquaculture* **364,** 165-172.

Kuukka-Anttila, H., Peuhkuri, N., Kolari, I., Paananen, T., Kause, A. (2010) Quantitative genetic architecture of parasite-induced cataract in rainbow trout, Oncorhynchus mykiss. *Heredity* **104,** 20-27.

Kvingedal, R., Evans, B.S., Lind, C.E., Taylor, J.J., Dupont-Nivet, M., Jerry, D.R. (2010) Population and family growth response to different rearing location, heritability estimates and genotype× environment interaction in the silver-lip pearl oyster (Pinctada maxima). *Aquaculture* **304,** 1-6.

Ky, C.-L., Blay, C., Sham-Koua, M., Lo, C., Cabral, P. (2014) Indirect improvement of pearl grade and shape in farmed Pinctada margaritifera by donor “oyster” selection for green pearls. *Aquaculture* **432,** 154-162.

LaFrentz, B.R., Lozano, C.A., Shoemaker, C.A., *et al.* (2016) Controlled challenge experiment demonstrates substantial additive genetic variation in resistance of Nile tilapia (Oreochromis niloticus) to Streptococcus iniae. *Aquaculture* **458,** 134-139.

Langdon, C., Evans, F., Jacobson, D., Blouin, M. (2003) Yields of cultured Pacific oysters Crassostrea gigas Thunberg improved after one generation of selection. *Aquaculture* **220,** 227-244.

Le Boucher, R., Quillet, E., Vandeputte, M.*, et al.* (2011) Plant-based diet in rainbow trout (Oncorhynchus mykiss Walbaum): Are there genotype-diet interactions for main production traits when fish are fed marine vs. plant-based diets from the first meal? *Aquaculture* **321,** 41-48.

Le Boucher, R., Vandeputte, M., Dupont-Nivet, M.*, et al.* (2013) Genotype by diet interactions in European sea bass (*Dicentrarchus labrax L.*): Nutritional challenge with totally plant-based diets. *Journal of Animal Science* **91,** 44-56.

Leaver, M.J., Taggart, J.B., Villeneuve, L.*, et al.* (2011) Heritability and mechanisms of n− 3 long chain polyunsaturated fatty acid deposition in the flesh of Atlantic salmon. *Comparative Biochemistry and Physiology Part D: Genomics and Proteomics* **6,** 62-69.

Leeds, T., Silverstein, J., Weber, G.*, et al.* (2010) Response to selection for bacterial cold water disease resistance in rainbow trout. *Journal of Animal Science* **88,** 1936-1946.

Leeds, T.D., Vallejo, R.L., Weber, G.M., Gonzalez-Pena, D., Silverstein, J.T. (2016) Response to five generations of selection for growth performance traits in rainbow trout (Oncorhynchus mykiss). *Aquaculture* **465,** 341-351.

Lhorente, J.P., Gallardo, J.A., Villanueva, B.*, et al.* (2012) Quantitative genetic basis for resistance to Caligus rogercresseyi sea lice in a breeding population of Atlantic salmon (Salmo salar). *Aquaculture* **324,** 55-59.

Lhorente, J.P., Gallardo, J.A., Villanueva, B., Carabaño, M.J., Neira, R. (2014) Disease resistance in Atlantic salmon (Salmo salar): coinfection of the intracellular bacterial pathogen Piscirickettsia salmonis and the sea louse Caligus rogercresseyi. *PloS One* **9,** e95397.

Li, Q., Wang, Q., Liu, S., Kong, L. (2011) Selection response and realized heritability for growth in three stocks of the Pacific oyster Crassostrea gigas. *Fisheries Science* **77,** 643-648.

Li, W., Luan, S., Luo, K.*, et al.* (2015) Genetic parameters and genotype by environment interaction for cold tolerance, body weight and survival of the Pacific white shrimp Penaeus vannamei at different temperatures. *Aquaculture* **441,** 8-15.

Liang, B., Jiang, F., Zhang, S., Yue, X., Wang, H., Liu, B. (2017) Genetic variation in vibrio resistance in the clam Meretrix petechialis under the challenge of Vibrio parahaemolyticus. *Aquaculture* **468,** 458-463.

Liang, J., Zhang, G., Zheng, H. (2010) Divergent selection and realized heritability for growth in the Japanese scallop, Patinopecten yessoensis Jay. *Aquaculture Research* **41,** 1315-1321.

Lillehammer, M., Ødegård, J., Madsen, P., Gjerde, B., Refstie, T., Rye, M. (2013) Survival, growth and sexual maturation in Atlantic salmon exposed to infectious pancreatic necrosis: a multi-variate mixture model approach. *Genetics, selection, evolution: GSE* **45,** 8.

Liu, B., Tan, C., Zhang, D.*, et al.* (2016) Genetic parameters of growth and resistance to Polydora ciliata in the pearl oyster Pinctada fucata. *Aquaculture Research*.

Liu, J., Lai, Z., Fu, X.*, et al.* (2015) Genetic parameters and selection responses for growth and survival of the small abalone Haliotis diversicolor after four generations of successive selection. *Aquaculture* **436,** 58-64.

Luan, S., Luo, K., Chai, Z.*, et al.* (2015a) An analysis of indirect genetic effects on adult body weight of the Pacific white shrimp Litopenaeus vannamei at low rearing density. *Genetics Selection Evolution* **47,** 95.

Luan, S., Wang, J., Yang, G.*, et al.* (2015b) Genetic parameters of survival for six generations in the giant freshwater prawn Macrobrachium rosenbergii. *Aquaculture Research* **46,** 1345-1355.

Luan, S., Yang, G., Wang, J.*, et al.* (2012) Genetic parameters and response to selection for harvest body weight of the giant freshwater prawn *Macrobrachium rosenbergii*. *Aquaculture* **262-263,** 88-96.

Luan, T.D., Olesen, I., Ødegård, J., Kolstad, K., Dan, N.C. (2008) Genotype by environment interaction for harvest body weight and survival of Nile tilapia (*Oreochromis niloticus*) in brackish and Fresh water ponds. In: *the proceedings of the 8th International Symposium on Tilapia in Aquaculture*. Egypt, pp. pp 231-238.

Lucas, T., Macbeth, M., Degnan, S.M., Knibb, W., Degnan, B.M. (2006) Heritability estimates for growth in the tropical abalone *Haliotis asinina* using microsatellites to assign parentage. *Aquaculture* **259,** 146-152.
Lyu, D., Wang, W., Luan, S., Hu, Y., Kong, J. (2017) Estimating genetic parameters for growth traits with molecular relatedness in turbot (Scophthalmus maximus, Linnaeus). *Aquaculture* **468,** 149-155.
Macbeth, M., Kenway, M., Salmon, M., Benzie, J., Knibb, W., Wilson, K. (2007) Heritability of reproductive traits and genetic correlations with growth in the black tiger prawn *Penaeus monodon* reared in tanks. *Aquaculture* **270,** 51-56.
Mahapatra, K.D., Gjerde, B., Sahoo, P.K*., et al.* (2008) Genetic variations in survival of rohu carp (*Labeo rohita, Hamilton*) after Aeromonas hydrophila infection in challenge tests. *Aquaculture* **279,** 29-34.
Maluwa, A.O., Gjerde, B. (2006) Estimates of the strain additive, maternal and heterosis genetic effects for harvest body weight of an F2 generation of *Oreochromis shiranus*. *Aquaculture* **259,** 38-46.
Maluwa, A.O., Gjerde, B., Ponzoni, R.W. (2006) Genetic parameters and genotype by environment interaction for body weight of *Oreochromis shiranus*. *Aquaculture* **259,** 47-55.
Marjanovic, J., Mulder, H.A., Khaw, H.L., Bijma, P. (2016) Genetic parameters for uniformity of harvest weight and body size traits in the GIFT strain of Nile tilapia. *Genetics Selection Evolution* **48,** 41.
Mas-Muñoz, J., Blonk, R., Schrama, J.W., van Arendonk, J., Komen, H. (2013) Genotype by environment interaction for growth of sole (*Solea solea*) reared in an intensive aquaculture system and in a semi-natural environment. *Aquaculture* **410–411,** 230-235.
McKay, L.R., Gjerde, B. (1986) Genetic variation for a spinal deformity in Atlantic salmon, Salmo salar. *Aquaculture* **52,** 263-272.
McPhee, C.P., Jones, C.M., Shanks, S.A. (2004) Selection for increased weight at 9 months in redclaw crayfish (*Cherax quadricarinatus*). *Aquaculture* **237,** 131-140.
Millot, S., Péan, S., Labbé, L*., et al.* (2014) Assessment of genetic variability of fish personality traits using rainbow trout isogenic lines. *Behavior genetics* **44,** 383-393.
Mohanty, B., Sahoo, P., Mahapatra, K., Saha, J. (2012) Differential resistance to edwardsiellosis in rohu (Labeo rohita) families selected previously for higher growth and/or aeromoniasis-resistance. *Journal of Applied Genetics* **53,** 107-114.
Montaldo, H.H., Castillo-Juárez, H., Campos-Montes, G., Pérez-Enciso, M. (2012) Effect of the data family structure, tank replication and the statistical model, on the estimation of genetic parameters for body weight at 28 days of age in the Pacific white shrimp (Penaeus (Litopenaeus) vannamei Boone, 1931). *Aquaculture Research***,** in press.
Moss, S.M., Moss, D.R., Arce, S.M., Lightner, D.V., Lotz, J.M. (2012) The role of selective breeding and biosecurity in the prevention of disease in penaeid shrimp aquaculture. *Journal of Invertebrate Pathology* **110,** 247-250.
Mustafa, A., MacKinnon, B. (1999) Genetic variation in susceptibility of Atlantic salmon to the sea louse Caligus elongatus Nordmann, 1832. *Canadian Journal of Zoology* **77,** 1332-1335.
Myers, J.M., Hershberger, W.K., Saxton, A.M., Iwamoto, R.N. (2001) Estimates of genetic and phenotypic parameters for length and weight of marine net-pen reared coho salmon (*Oncorhynchus kisutch Walbaum*). *Aquaculture Research* **32,** 277-285.
Navarro, A., Zamorano, M.J., Hildebrandt, S., Ginés, R., Aguilera, C., Afonso, J.M. (2009a) Estimates of heritabilities and genetic correlations for body composition traits and G × E interactions, in gilthead seabream (*Sparus auratus L.*). *Aquaculture* **295,** 183-187.
Navarro, A., Zamorano, M.J., Hildebrandt, S., Ginés, R., Aguilera, C., Afonso, J.M. (2009b) Estimates of heritabilities and genetic correlations for growth and carcass traits in gilthead seabream (*Sparus auratus L.*), under industrial conditions. *Aquaculture* **289,** 225-230.

Neira, R., Díaz, N.F., Gall, G.A., Gallardo, J.A., Lhorente, J.P., Alert, A. (2006) Genetic improvement in coho salmon (Oncorhynchus kisutch). II: Selection response for early spawning date. *Aquaculture* **257,** 1-8.

Neira, R., Lhorente, J.P., Araneda, C., Díaz, N., Bustos, E., Alert, A. (2004) Studies on carcass quality traits in two populations of Coho salmon (*Oncorhynchus kisutch*): phenotypic and genetic parameters. *Aquaculture* **241,** 117-131.

Nell, J.A., Perkins, B. (2005) Evaluation of progeny of fourth generation Sydney rock oyster Saccostrea glomerata (Gould, 1850) breeding lines. *Aquaculture Research* **36,** 753-757.

Nguyen, N.H. (2015) Genetic improvement for important farmed aquaculture species with a reference to carp, tilapia and prawns in Asia: achievements, lessons and challenges. *Fish and Fisheries***,** doi: 10.1111/faf.12122.

Nguyen, N.H., Khaw, H.L., Ponzoni, R.W., Hamzah, A., Kamaruzzaman, N. (2007) Can sexual dimorphism and body shape be altered in Nile tilapia (*Oreochromis niloticus*) by genetic means? *Aquaculture* **272,** S38-S46.

Nguyen, N.H., Ponzoni, R.W., Abu-Bakar, K.R., Hamzah, A., Khaw, H.L., Yee, H.Y. (2010a) Correlated response in fillet weight and yield to selection for increased harvest weight in genetically improved farmed tilapia (GIFT strain), *Oreochromis niloticus*. *Aquaculture* **305,** 1-5.

Nguyen, N.H., Ponzoni, R.W., Yee, H.Y., Abu-Bakar, K.R., Hamzah, A., Khaw, H.L. (2010b) Quantitative genetic basis of fatty acid composition in the GIFT strain of Nile tilapia (*Oreochromis niloticus*) selected for high growth. *Aquaculture* **309,** 66-74.

Nguyen, N.H., Quinn, J., Powell, D.*, et al.* (2014a) Heritability for body colour and its genetic association with morphometric traits in Banana shrimp (Fenneropenaeus merguiensis). *BMC Genetics* **15,** 132.

Nguyen, N.H., Whatmore, P., Miller, A., Knibb, W. (2015) Quantitative genetic properties of four measures of deformity in yellowtail kingfish Seriola lalandi Valenciennes, 1833. *Journal of Fish Diseases***,** doi: 10.1111/jfd.12348.

Nguyen, T., Hayes, B., Guthridge, K., Ab Rahim, E., Ingram, B. (2011) Use of a microsatellite-based pedigree in estimation of heritabilities for economic traits in Australian blue mussel, Mytilus galloprovincialis. *Journal of Animal Breeding and Genetics* **128,** 482-490.

Nguyen, T.T., Hayes, B.J., Ingram, B.A. (2014b) Genetic parameters and response to selection in blue mussel (Mytilus galloprovincialis) using a SNP-based pedigree. *Aquaculture* **420,** 295-301.

Nielsen, H.M., Monsen, B.B., Ødegård, J.*, et al.* (2014) Direct and social genetic parameters for growth and fin damage traits in Atlantic cod (Gadus morhua). *Genet Sel Evol* **46**.

Nielsen, H.M., Ødegård, J., Olesen, I.*, et al.* (2010) Genetic analysis of common carp (*Cyprinus carpio*) strains: I: Genetic parameters and heterosis for growth traits and survival. *Aquaculture* **304,** 14-21.

Nilsson, J., Backström, T., Stien, L.*, et al.* (2016) Effects of age and rearing environment on genetic parameters of growth and body weight and heritability of skin pigmentation in Arctic charr (Salvelinus alpinus L.). *Aquaculture* **453,** 67-72.

Nilsson, J., Brännäs, E., Eriksson, L.-O. (2010) The Swedish Arctic charr breeding programme. *Hydrobiologia* **650,** 275-282.

Ninh, N.H., Ponzoni, R.W., Nguyen, N.H., Woolliams, J.A., McAndrew, B.J., Penman, D.J. (2011) Communal or separate rearing of families in selective breeding of common carp (*Cyprinus carpio*). *Aquaculture* **in press**.

Ninh, N.H., Thoa, N.P., Knibb, W., Nguyen, N.H. (2014) Selection for enhanced growth performance of Nile tilapia (*Oreochromis niloticus*) in brackish water (15 - 20 ppt) in Vietnam. *Aquaculture* **428-429,** 1-6.

Norris, A., Cunningham, E. (2004) Estimates of phenotypic and genetic parameters for flesh colour traits in farmed Atlantic salmon based on multiple trait animal model. *Livestock Production Science* **89,** 209-222.

Norris, A., Foyle, L., Ratcliff, J. (2008) Heritability of mortality in response to a natural pancreas disease (SPDV) challenge in Atlantic salmon, Salmo salar L., post-smolts on a West of Ireland sea site. *Journal of Fish Diseases* **31,** 913-920.
Ødegård, J., Baranski, M., Gjerde, B., Gjedrem, T. (2011a) Methodology for genetic evaluation of disease resistance in aquaculture species: challenges and future prospects. *Aquaculture Research* **42,** 103-114.
Ødegård, J., Gitterle, T., Madsen, P.*, et al.* (2011b) Quantitative genetics of taura syndrome resistance in pacific white shrimp (penaeus vannamei): a cure model approach. *Genetics Selection Evolution* **43,** 14.
Ødegård, J., Olesen, I., Dixon, P.*, et al.* (2010) Genetic analysis of common carp (*Cyprinus carpio*) strains. II: Resistance to koi herpesvirus and Aeromonas hydrophila and their relationship with pond survival. *Aquaculture* **304,** 7-13.
Ødegård, J., Olesen, I., Gjerde, B., Klemetsdal, G. (2007) Positive genetic correlation between resistance to bacterial (furunculosis) and viral (infectious salmon anaemia) diseases in farmed Atlantic salmon (Salmo salar). *Aquaculture* **271,** 173-177.
Oldorf, W., Kronert, U., Balarin, J., Haller, R., Hörstgen-Schwark, G., Langholz, H.-J. (1989) Prospects of selecting for late maturity in tilapia (Oreochromis niloticus): II. Strain comparisons under laboratory and field conditions. *Aquaculture* **77,** 123-133.
Olesen, I., Hung, D., Ødegård, J. (2007) Genetic analysis of survival in challenge tests of furunculosis and ISA in Atlantic salmon. Genetic parameter estimates and model comparisons. *Aquaculture* **272,** S297-S298.
Oliveira, C.A.L., Ribeiro, R.P., Yoshida, G.M.*, et al.* (2016) Correlated changes in body shape after five generations of selection to improve growth rate in a breeding program for Nile tilapia Oreochromis niloticus in Brazil. *Journal of Applied Genetics*, 1-7. DOI 10.1007/s13353-13016-10338-13355.
Omasaki, S., Charo-Karisa, H., Kahi, A., Komen, H. (2016) Genotype by environment interaction for harvest weight, growth rate and shape between monosex and mixed sex Nile tilapia (Oreochromis niloticus). *Aquaculture* **458,** 75-81.
Pante, M.J.R., Gjerde, B., McMillan, I., Misztal, I. (2002) Estimation of additive and dominance genetic variances for body weight at harvest in rainbow trout, *Oncorhynchus mykiss*. *Aquaculture* **204,** 383-392.
Pérez-Rostro, C.I., Ibarra, A.M. (2003) Heritabilities and genetic correlations of size traits at harvest size in sexually dimorphic Pacific white shrimp (*Litopenaeus vannamei*) grown in two environments. *Aquaculture Research* **34,** 1079-1085.
Pérez-Rostro, C.I., Ramirez, J.L., Ibarra, A.M. (1999) Maternal and cage effects on genetic parameter estimation for Pacific white shrimp *Penaeus vannamei* Boone. *Aquaculture Research* **30,** 681-693.
Perez, J., Alfonsi, C. (1999) Selection and realized heritability for growth in the scallop, Euvola ziczac (L.). *Aquaculture Research* **30,** 211-214.
Perry, G.M.L., Martyniuk, C.M., Ferguson, M.M., Danzmann, R.G. (2005) Genetic parameters for upper thermal tolerance and growth-related traits in rainbow trout (*Oncorhynchus mykiss*). *Aquaculture* **250,** 120-128.
Perry, G.M.L., Tarte, P., Croisetière, S., Belhumeur, P., Bernatchez, L. (2004) Genetic variance and covariance for 0+ brook charr (Salvelinus fontinalis) weight and survival time of furunculosis (Aeromonas salmonicida) exposure. *Aquaculture* **235,** 263-271.
Phuthaworn, C., Nguyen, N.H., Quinn, J., Knibb, W. (2016) Moderate heritability of hepatopancreatic parvovirus titre suggests a new option for selection against viral diseases in banana shrimp (Fenneropenaeus merguiensis) and other aquaculture species. *Genetics Selection Evolution* **48,** 64.

Pierce, L.R., Palti, Y., Silverstein, J.T., Barrows, F.T., Hallerman, E.M., Parsons, J.E. (2008) Family growth response to fishmeal and plant-based diets shows genotype× diet interaction in rainbow trout (Oncorhynchus mykiss). *Aquaculture* **278,** 37-42.

Pino-Querido, A., Álvarez-Castro, J.M., Guerra-Varela, J*., et al.* (2015) Heritability estimation for okadaic acid algal toxin accumulation, mantle color and growth traits in Mediterranean mussel (Mytilus galloprovincialis). *Aquaculture* **440,** 32-39.

Ponzoni, R.W., Hamzah, A., Tan, S., Kamaruzzaman, N. (2005) Genetic parameters and response to selection for live weight in the GIFT strain of Nile Tilapia (*Oreochromis niloticus*). *Aquaculture* **247,** 203-210.

Powell, J., White, I., Guy, D., Brotherstone, S. (2008) Genetic parameters of production traits in Atlantic salmon (*Salmo salar*). *Aquaculture* **274,** 225-231.

Quillet, E., Le Guillou, S., Aubin, J., Fauconneau, B. (2005) Two-way selection for muscle lipid content in pan-size rainbow trout (Oncorhynchus mykiss). *Aquaculture* **245,** 49-61.

Quillet, E., Le Guillou, S., Aubin, J., Labbé, L., Fauconneau, B., Médale, F. (2007) Response of a lean muscle and a fat muscle rainbow trout (Oncorhynchus mykiss) line on growth, nutrient utilization, body composition and carcass traits when fed two different diets. *Aquaculture* **269,** 220-231.

Quinton, C.D., Kause, A., Ruohonen, K., Koskela, J. (2007) Genetic relationships of body composition and feed utilization traits in European whitefish (*Coregonus lavaretus L.*) and implications for selective breeding in fishmeal- and soybean meal-based diet environments. *Journal of Animal Science* **85,** 3198-3208.

Quinton, C.D., McMillan, I., Glebe, B.D. (2005) Development of an Atlantic salmon (*Salmo salar*) genetic improvement program: Genetic parameters of harvest body weight and carcass quality traits estimated with animal models. *Aquaculture* **247,** 211-217.

Rezk, M.A., Ponzoni, R.W., Khaw, H.L., Kamel, E., Dawood, T., John, G. (2009) Selective breeding for increased body weight in a synthetic breed of Egyptian Nile tilapia, *Oreochromis niloticus*: Response to selection and genetic parameters. *Aquaculture* **293,** 187-194.

Rutten, M.J.M. (2005) Breeding for improved production of Nile Tilapia (*Oreochromis niloticus* L.). Doctorial thesis, Doctorial thesis. Animal Breeding and Genetic Group, Wageningen University.

Rye, M., Gjerde, B. (1996) Phenotypic and genetic parameters of body composition traits and flesh colour in Atlantic salmon, Salmo salar L. *Aquaculture Research* **27,** 121-133.

Rye, M., Lillevik, K.M. (1990) Survival in the early fresh-water period in Atlantic salmon (*Salmo salar*) and rainbow trout (*Salmo gairdneri*): heritabilities for survival and genetic correlation between survival and growth. *Aquaculture* **85,** 328-329.

Rye, M., Mao, I.L. (1998) Nonadditive genetic effects and inbreeding depression for body weight in Atlantic salmon (*Salmo salar* L.). *Livestock Production Science* **57,** 15-22.

Sae-Lim, P., Kause, A., Janhunen, M*., et al.* (2015) Genetic (co) variance of rainbow trout (Oncorhynchus mykiss) body weight and its uniformity across production environments. *Genetics Selection Evolution* **47,** 1-10.

Sae-Lim, P., Kause, A., Mulder, H.A*., et al.* (2013) Genotype-by-environment interaction of growth traits in rainbow trout (*Oncorhynchus mykiss*): A continental scale study. *Journal of Animal Science* **91,** 5572-5581.

Sahoo, B.R., Basu, M., Swain, B., Dikhit, M.R., Jayasankar, P., Samanta, M. (2013) Elucidation of novel structural scaffold in rohu TLR2 and its binding site analysis with peptidoglycan, lipoteichoic acid and zymosan ligands, and downstream MyD88 adaptor protein. *BioMed research international* **2013**.

Sahoo, P., Mahapatra, K.D., Saha, J*., et al.* (2008) Family association between immune parameters and resistance to Aeromonas hydrophila infection in the Indian major carp, Labeo rohita. *Fish & Shellfish Immunology* **25,** 163-169.

Saillant, E., Dupont-Nivet, M., Sabourault, M., *et al.* (2009) Genetic variation for carcass quality traits in cultured sea bass ( Dicentrarchus labrax). *Aquatic Living Resources* **22,** 105-112.
Salte, R., Bentsen, H.B., Moen, T., *et al.* (2009) Prospects for a genetic management strategy to control Gyrodactylus salaris infection in wild Atlantic salmon (Salmo salar) stocks. *Canadian Journal of Fisheries and Aquatic Sciences* **67,** 121-129.
Samain, J.-F., Degremont, L., Soletchnik, P., *et al.* (2007) Genetically based resistance to summer mortality in the Pacific oyster (Crassostrea gigas) and its relationship with physiological, immunological characteristics and infection processes. *Aquaculture* **268,** 227-243.
Sang, N.V., Klemetsdal, G., Odegard, J., Gjoen, H.M. (2012) Genetic parameters of economically important traits recorded at a given age in striped catfish (*Pangasianodon hypophthalmus*). *Aquaculture* **344 - 349,** 82-89.
Sheridan, A. (1997) Genetic improvement of oyster production—a critique. *Aquaculture* **153,** 165-179.
Shoemaker, C.A., Lozano, C.A., LaFrentz, B.R., *et al.* (2017) Additive genetic variation in resistance of Nile tilapia (Oreochromis niloticus) to Streptococcus iniae and S. agalactiae capsular type Ib: Is genetic resistance correlated? *Aquaculture* **468,** 193-198.
Silverstein, J., Vallejo, R., Palti, Y., *et al.* (2009) Rainbow trout resistance to bacterial cold-water disease is moderately heritable and is not adversely correlated with growth. *Journal of Animal Science* **87,** 860-867.
Silverstein, J.T., Bosworth, B.G., Waldbieser, G.C., Wolters, W.R. (2001) Feed intake in channel catfish: is there a genetic component? *Aquaculture Research* **32,** 199-205.
Sonesson, A.K., Ødegård, J., Rönnegård, L. (2013) Genetic heterogeneity of within-family variance of body weight in Atlantic salmon (Salmo salar). *Genetics Selection Evolution* **45,** 41.
Su, G.-S., Liljedahl, L.-E., Gall, G.A. (1996) Genetic and environmental variation of body weight in rainbow trout (Oncorhynchus mykiss). *Aquaculture* **144,** 71-80.
Su, G.-S., Liljedahl, L.-E., Gall, G.A. (1997) Genetic and environmental variation of female reproductive traits in rainbow trout (Oncorhynchus mykiss). *Aquaculture* **154,** 115-124.
Su, G.-S., Liljedahl, L.-E., Gall, G.A. (2002a) Genetic correlations between body weight at different ages and with reproductive traits in rainbow trout. *Aquaculture* **213,** 85-94.
Su, G.-S., Liljedahl, L.-E., Gall, G.A.E. (2002b) Genetic correlations between body weight at different ages and with reproductive traits in rainbow trout. *Aquaculture* **213,** 85-94.
Sui, J., Luan, S., Luo, K., *et al.* (2015) Genetic parameters and response to selection for harvest body weight of pacific white shrimp, Litopenaeus vannamei. *Aquaculture Research*, n/a-n/a.
Sun, M.M., Huang, J.H., Jiang, S.G., *et al.* (2015) Estimates of heritability and genetic correlations for growth-related traits in the tiger prawn Penaeus monodon. *Aquaculture Research* **46,** 1363-1368.
Swan, A.A., Thompson, P.A., Ward, R.D. (2007a) Genotype x environment interactions for weight in Pacific oysters (*Crassostrea gigas*) on five Australian farms. *Aquaculture* **265,** 91-101.
Swan, A.A., Thompson, P.A., Ward, R.D. (2007b) Genotype × environment interactions for weight in Pacific oysters (*Crassostrea gigas*) on five Australian farms. *Aquaculture* **265,** 91-101.
Sylvén, S., Rye, M., Simianer, H. (1991) Interaction of genotype with production system for slaughter weight in rainbow trout (*Oncorhynchus mykiss*). *Livestock Production Science* **28,** 253-263.
Tan, J., Kong, J., Cao, B., *et al.* (2017) Genetic parameter estimation of reproductive traits of Litopenaeus vannamei. *Journal of Ocean University of China* **16,** 161-167.
Taylor, R.S., Muller, W.J., Cook, M.T., Kube, P.D., Elliott, N.G. (2009) Gill observations in Atlantic salmon (Salmo salar, L.) during repeated amoebic gill disease (AGD) field exposure and survival challenge. *Aquaculture* **290,** 1-8.
Thoa, N.P., Knibb, W., Ninh, N.H., *et al.* (2015) Genetic variation in survival of tilapia (Oreochromis niloticus, Linnaeus, 1758) fry during the early phase of rearing in brackish water environment (5–10 ppt). *Aquaculture* **442,** 112-118.

Thoa, N.P., Ninh, N.H., Knibb, W., Nguyen, N.H. (2016) Does selection in a challenging environment produce Nile tilapia genotypes that can thrive in a range of production systems? *Scientific Reports* **6,** 21486 doi:21410.21038/srep21486.

Thodesen, J., Rye, M., Wang, Y.-X., Li, S.-J., Bentsen, H.B., Gjedrem, T. (2013a) Genetic improvement of tilapias in China: Genetic parameters and selection responses in growth, pond survival and cold-water tolerance of blue tilapia (Oreochromis aureus) after four generations of multi-trait selection. *Aquaculture* **396–399,** 32-42.

Thodesen, J., Rye, M., Wang, Y.-X.*, et al.* (2013b) Genetic improvement of tilapias in China: Genetic parameters and selection responses in growth, survival and external color traits of red tilapia (Oreochromis spp.) after four generations of multi-trait selection. *Aquaculture* **416,** 354-366.

Tobin, D., Kause, A., Mäntysaari, E.A.*, et al.* (2006) Fat or lean? The quantitative genetic basis for selection strategies of muscle and body composition traits in breeding schemes of rainbow trout (Oncorhynchus mykiss). *Aquaculture* **261,** 510-521.

Toro, J.E., Alcapán, A.C., Vergara, A.M., Ojeda, J.A. (2004) Heritability estimates of larval and spat shell height in the Chilean blue mussel (Mytilus chilensis Hupe 1854) produced under controlled laboratory conditions. *Aquaculture Research* **35,** 56-61.

Toro, J.E., Newkirk, G.F. (1991) Response to artificial selection and realized heritability estimate for shell height in the Chilean oyster Ostrea chilensis. *Aquatic Living Resources* **4,** 101-108.

Tosh, J., Garber, A., Trippel, E., Robinson, J. (2010) Genetic, maternal, and environmental variance components for body weight and length of Atlantic cod at 2 points in life. *Journal of Animal Science* **88,** 3513-3521.

Trọng, T.Q., Mulder, H.A., van Arendonk, J.A.M., Komen, H. (2013a) Heritability and genotype by environment interaction estimates for harvest weight, growth rate, and shape of Nile tilapia (*Oreochromis niloticus*) grown in river cage and VAC in Vietnam. *Aquaculture* **384–387,** 119-127.

Trọng, T.Q., van Arendonk, J.A., Komen, H. (2013b) Genetic parameters for reproductive traits in female Nile tilapia (Oreochromis niloticus): I. Spawning success and time to spawn. *Aquaculture* **416,** 57-64.

Trọng, T.Q., van Arendonk, J.A., Komen, H. (2013c) Genetic parameters for reproductive traits in female Nile tilapia (Oreochromis niloticus): II. Fecundity and fertility. *Aquaculture* **416,** 72-77.

Trọng, T.Q., van Bers, N., Crooijmans, R., Dibbits, B., Komen, H. (2013d) A comparison of microsatellites and SNPs in parental assignment in the GIFT strain of Nile tilapia (*Oreochromis niloticus*): The power of exclusion. *Aquaculture* **388,** 14-23.

Turra, E.M., Toral, F.L.B., de Alvarenga, É.R.*, et al.* (2016) Genotype× environment interaction for growth traits of Nile tilapia in biofloc technology, recirculating water and Cage systems. *Aquaculture* **460,** 98-104.

Vallejo, R.L., Leeds, T.D., Gao, G.*, et al.* (2017) Genomic selection models double the accuracy of predicted breeding values for bacterial cold water disease resistance compared to a traditional pedigree-based model in rainbow trout aquaculture. *Genetics Selection Evolution* **49,** 17.

Vandeputte, M. (2003) Selective breeding of quantitative traits in the common carp (*Cyprinus carpio*): a review. *Aquatic Living Resources* **16,** 399-407.

Vandeputte, M., Kocour, M., Mauger, S.*, et al.* (2004) Heritability estimates for growth-related traits using microsatellite parentage assignment in juvenile common carp (*Cyprinus carpio L.*). *Aquaculture* **235,** 223-236.

Vandeputte, M., Kocour, M., Mauger, S.*, et al.* (2008) Genetic variation for growth at one and two summers of age in the common carp (*Cyprinus carpio* L.): Heritability estimates and response to selection. *Aquaculture* **277,** 7-13.

Vandeputte, M., Porte, J., Auperin, B.*, et al.* (2016) Quantitative genetic variation for post-stress cortisol and swimming performance in growth-selected and control populations of European sea bass (Dicentrarchus labrax). *Aquaculture* **455,** 1-7.
Vandeputte, M., Puledda, A., Tyran, A.S.*, et al.* (2017) Investigation of morphological predictors of fillet and carcass yield in European sea bass (Dicentrarchus labrax) for application in selective breeding. *Aquaculture* **470,** 40-49.
Vandeputte, M., Quillet, E., Chevassus, B. (2002) Early development and survival in brown trout (*Salmo trutta fario L.*): indirect effects of selection for growth rate and estimation of genetic parameters. *Aquaculture* **204,** 435-445.
Vehviläinen, H., Kause, A., Koskinen, H., Paananen, T. (2010) Genetic architecture of rainbow trout survival from egg to adult. *Genetics research* **92,** 1-11.
Vehviläinen, H., Kause, A., Kuukka-Anttila, H., Koskinen, H., Paananen, T. (2012) Untangling the positive genetic correlation between rainbow trout growth and survival. *Evolutionary applications* **5,** 732-745.
Vehviläinen, H., Kause, A., Quinton, C., Koskinen, H., Paananen, T. (2008) Survival of the currently fittest: genetics of rainbow trout survival across time and space. *Genetics* **180,** 507-516.
Vieira, V.L.A., Norris, A., Johnston, I.A. (2007) Heritability of fibre number and size parameters and their genetic relationship to flesh quality traits in Atlantic salmon (*Salmo salar L.*). *Aquaculture* **272,** S100-S109.
Volckaert, F.A., Hellemans, B., Batargias, C.*, et al.* (2012) Heritability of cortisol response to confinement stress in European sea bass dicentrarchus labrax. *Genetics Selection Evolution* **44,** 15.
Wada, K.T., Komaru, A. (1994) Effect of selection for shell coloration on growth rate and mortality in the Japanese pearl oyster, Pinctada fucata martensii. *Aquaculture* **125,** 59-65.
Wang, C.-h., Li, S.-f., Xiang, S.-p.*, et al.* (2006a) Genetic parameter estimates for growth-related traits in Oujiang color common carp (*Cyprinus carpio* var. color). *Aquaculture* **259,** 103-107.
Wang, C.M., Lo, L.C., Zhu, Z.Y.*, et al.* (2008) Estimating reproductive success of brooders and heritability of growth traits in Asian sea bass (Lates calcarifer) using microsatellites. *Aquaculture Research* **39,** 1612-1619.
Wang, H., Du, X., Lü, W., Liu, Z. (2010) Estimating the heritability for growth-related traits in the pearl oyster, Pinctada fucata martensii (Dunker). *Aquaculture Research* **42,** 57-64.
Wang, X., Ross, K.E., Saillant, E., Gatlin Iii, D.M., Gold, J.R. (2006b) Quantitative genetics and heritability of growth-related traits in hybrid striped bass (*Morone chrysops* [female symbol] × *Morone saxatilis* [male symbol]). *Aquaculture* **261,** 535-545.
Wetten, M., Aasmundstad, T., Kjøglum, S., Storset, A. (2007) Genetic analysis of resistance to infectious pancreatic necrosis in Atlantic salmon (Salmo salar L.). *Aquaculture* **272,** 111-117.
Whatmore, P., Nguyen, N.H., Miller, A.*, et al.* (2013) Genetic parameters for economically important traits in yellowtail kingfish *Seriola lalandi*. *Aquaculture* **400–401,** 77-84.
Wild, V., Simianer, H., Gjøen, H.M., Gjerde, B. (1994) Genetic parameters and genotype × environment interaction for early sexual maturity in Atlantic salmon (*Salmo salar*). *Aquaculture* **128,** 51-65.
Winkelman, A., Peterson, R. (1994a) Heritabilities, dominance variation, common environmental effects and genotype by environment interactions for weight and length in chinook salmon. *Aquaculture* **125,** 17-30.
Winkelman, A.M., Peterson, R.G. (1994b) Genetic parameters (heritabilities, dominance ratios and genetic correlations) for body weight and length of chinook salmon after 9 and 22 months of saltwater rearing. *Aquaculture* **125,** 31-36.
Withler, R., Beacham, T. (1994) Genetic variation in body weight and flesh color of the coho salmon (Oncorhynchus kisutch) in British Columbia. *Aquaculture* **119,** 135 - 148.
Yan, X., Huo, Z., Yang, F., Zhang, G. (2014) Heritability of larval and juvenile growth for two stocks of Manila clam Ruditapes philippinarum. *Aquaculture Research* **45,** 484-490.

Yáñez, J.M., Bangera, R., Lhorente, J.P., *et al.* (2016) Negative genetic correlation between resistance against Piscirickettsia salmonis and harvest weight in coho salmon (Oncorhynchus kisutch). *Aquaculture* **459,** 8-13.

Yáñez, J.M., Bangera, R., Lhorente, J.P., Oyarzún, M., Neira, R. (2013) Quantitative genetic variation of resistance against Piscirickettsia salmonis in Atlantic salmon (Salmo salar). *Aquaculture* **414,** 155-159.

Yáñez, J.M., Lhorente, J.P., Bassini, L.N., Oyarzún, M., Neira, R., Newman, S. (2014) Genetic co-variation between resistance against both Caligus rogercresseyi and Piscirickettsia salmonis, and body weight in Atlantic salmon (Salmo salar). *Aquaculture* **433,** 295-298.

Zak, T., Deshev, R., Benet-Perlberg, A., *et al.* (2014) Genetic improvement of Israeli blue (Jordan) tilapia, Oreochromis aureus (Steindachner), through selective breeding for harvest weight. *Aquaculture Research* **45,** 546-557.

Zhang, J., Cao, F., Liu, J., Yuan, R., Hu, Z. (2017) Genetic Parameters for Growth and Hypoxic Tolerance Traits in Pacific White Shrimp Litopenaeus vannamei at Different Ages. *North American Journal of Aquaculture* **79,** 75-83.

Zhao, H., Zeng, C., Wan, S., Dong, Z., Gao, Z. (2016) Estimates of Heritabilities and Genetic Correlations for Growth and Gonad Traits in Blunt Snout Bream, Megalobrama amblycephala. *Journal of the World Aquaculture Society* **47,** 139-146.

ASSESSMENT OF PUBLICATION BIAS

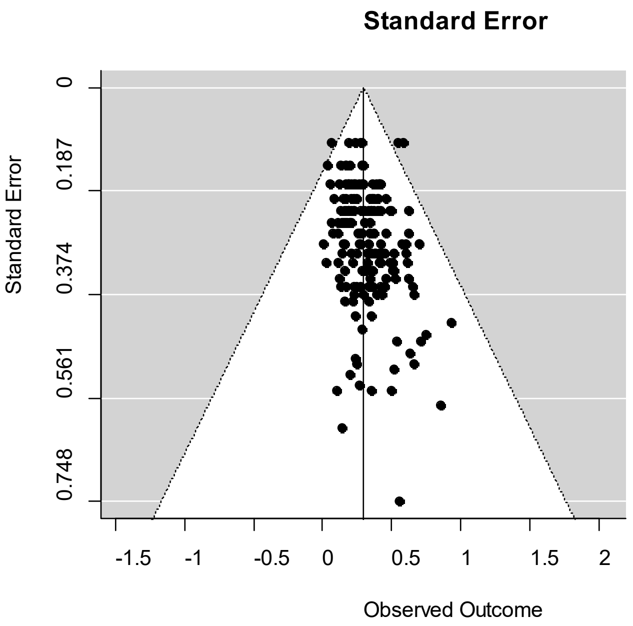

Funnel plot for body weight. The Egger test for funnel plot asymmetry: z = 1.950, p = 0.052

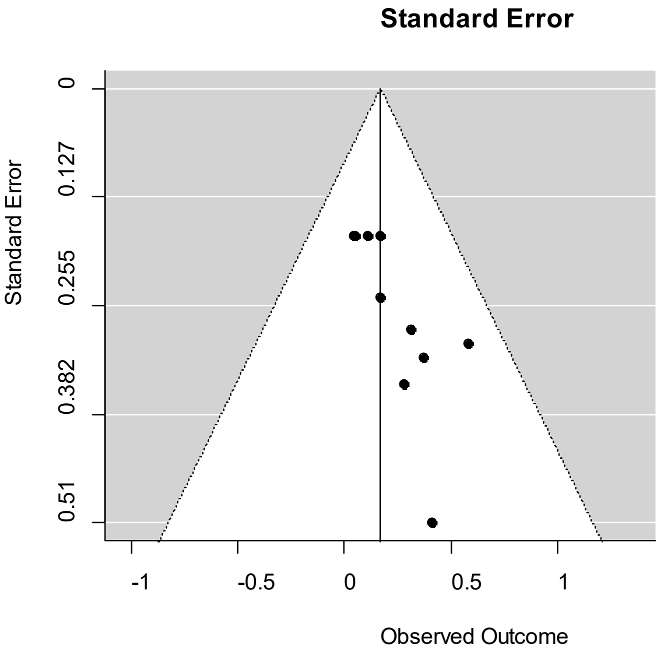

Funnel plot for fat content. The Egger test for funnel plot asymmetry: z = 1.559, p = 0.119

**Supplementary file N.4**: Description of novel traits discussed in this study

| Category | Trait name | Description | Method | References |
|---|---|---|---|---|
| Immune related parameters | | | | |
| | Cortisol | Cortisol was measured by radioimmunoassay | Laboratory analysis | Volckaert *et al.* 2012 |
| | Hypoxia stress | Hypoxia tolerance was recorded under challenge condition (DO < 2.0 mg/L) over consecutive days | Challenge test | Zhang *et al.* 2017 |
| Behaviour | | | | |
| | Risk-taking or boldness | The individual time-lapse to the first passage into the risky chamber; the total number of passages through the opaque divider for each individual; and risk-taking status was also referred as proactive, and risk avoiders | Video recording | Ferrari *et al.* 2016 |
| | Exploratory behaviour | Fish behaviour was recorded during 1 h after the stimulation (non-swimming fish characterized as "shy" and homogeneous swimming in the tank "bold" | Video recording | Millot *et al.* 2014 |
| | Behavioural responsiveness | time spent in central zone of the tank after the stress period (observed for 5 min after 2 min of stress), | Video recording | Drangsholt *et al.* 2014 |
| | Swimming behaviour | the swimming speed in body length per second (BL $s^{-1}$) after the stress period (observed for 5 min after 2 min of stress) | Video recording | Drangsholt *et al.* 2014 |
| Social interaction | Indirect genetic effect | Socially indirect genetic effects (IGEs) were estimated by extending the mixed model with IGEs | Group testing in the field | Bijma 2014 |
| Uniformity | Uniformity | Uniformity was estimated using double hierarchical linear generalised model in pedigreed populations | Pedigreed data | Rönnegård & Lee 2013 |
